# Uncovering microbial life-history strategies under disturbance: a trait-based computational analysis of anaerobic systems

**DOI:** 10.1101/2025.09.18.676720

**Authors:** Soheil A. Neshat, Ezequiel Santillan, Stefan Wuertz

## Abstract

Trait-based approaches are helpful in simplifying ecosystem complexity to explore disturbance-diversity-function relationships. These frameworks classify organisms based on their functional characteristics—traits that influence growth, survival and reproduction—providing a mechanistic basis to understand how communities respond to changes in their environment. The application of these approaches has been successful in ecology, but to date has only been tested in a few microbial ecosystems, namely, soil microbial communities and aerobic bioreactors treating wastewater. Here, we employed Grime’s competitor–stress-tolerant–ruderal framework in replicated mesophilic anaerobic bioreactors exposed to a disturbance (biomass removal) with varied frequencies at a constant number of disturbance events for 90 days. Bioreactors were inoculated with sludge from full-scale anaerobic digesters and fed with a mixture of primary and waste activated sludge. A genome-resolved metagenomics approach was utilised to assess the microbial communities. We found that communities across the disturbance range were clustered into three groups, suggesting the adoption of a three-way life-history strategy. This study demonstrates, for the first time, the applicability of trait-based life-history strategies in anaerobic microbial systems under disturbance, providing a new perspective for understanding and managing microbial ecosystems under disturbance conditions.

## 1. Introduction

Microbially mediated processes, driven by complex and dynamic microbial communities, are fundamental to modern wastewater treatment. The operational performance of these processes is directly linked to microbial community structure, which is often disrupted by environmental and operational disturbances (Santillan, Seshan, Constancias, Drautz-Moses, *et al*., 2019). While taxonomic profiling offers some insights into these communities, it lacks sufficient predictive power. Trait-based frameworks, which link community structure to ecosystem functions, offer a promising alternative for understanding and managing these complex systems (Krohn *et al*., 2022; Santillan, Neshat and Wuertz, 2025a).

Trait-based frameworks reduce ecological complexity by grouping organisms according to functional traits—attributes that influence fitness-related processes such as growth, reproduction, and survival. These frameworks typically map organisms onto conceptual axes that represent ecological trade-offs. Two-dimensional models, such as r/K selection (Pianka, 1970), generalists vs. specialists (Székely, Berga and Langenheder, 2013; Liao *et al*., 2016) and copiotrophy vs. oligotrophy (Lauro *et al*., 2009; Chen *et al*., 2021) have been widely used to study systems comprised of large organisms such as marine and land animals. However, such models often fail to capture the ecological nuances of microbial communities, which exhibit unique features such as dormancy, horizontal gene transfer, and rapid adaptation (Ho *et al*., 2013). To address such limitations, Ho et al. (Ho *et al*., 2013) proposed the adoption of Grime’s competitor–stress-tolerant–ruderal (CSR) framework, a three-dimensional trait-based model originally developed for plant ecology. This framework defines life-history strategies along three axes representing key ecological pressures: competition, stress, and disturbance. Its multidimensional structure offers a more flexible and ecologically meaningful way to characterize microbial strategies in dynamic environments, making it increasingly relevant for microbial ecology (Santillan, Neshat and Wuertz, 2025a). The application of such frameworks in microbial systems is often operationalized through Community Aggregated Traits (CATs), which summarize the distribution of traits across all community members. CATs offer a powerful means of linking metagenomic data to ecological strategy, enabling community-level positioning within trait spaces such as the CSR triangle (Santillan, Seshan, Constancias and Wuertz, 2019; Piton *et al*., 2023).

In the CSR model, organisms are classified into three primary life-history strategies: competitors (C), which maximize resource acquisition and control; stress-tolerants (S), which maintain metabolic activity under high-stress conditions; and ruderals (R), which exhibit rapid growth but low resource-use efficiency (Grime, 1977). C-strategists tend to dominate productive, stable environments, S-strategists thrive in unproductive or harsh conditions, and R-strategists are favoured in frequently disturbed environments. The balance among these strategies is shaped by three ecological forces: biomass destruction (disturbance), biomass restriction (stress), and resource competition. Meta-analyses have supported the applicability of the CSR framework in microbial ecology, particularly in soil ecosystem (Chagnon *et al*., 2013; Ho *et al*., 2013; Crowther *et al*., 2014; Krause, 2014; Ho, Di Lonardo and Bodelier, 2017; Piton *et al*., 2023).

Although promising, the CSR framework is still relatively new in the context of engineered microbial systems. To date, it has only been applied to activated sludge bioreactors treating synthetic wastewater under disturbance conditions (Santillan, Seshan, Constancias and Wuertz, 2019; Santillan, Neshat and Wuertz, 2025b). Such studies established an important precedent, demonstrating that a three-way trait-based framework can effectively link microbial community structure to function using metagenomics in controlled microcosm experiments. These studies also introduced and tested the hypothesis that undisturbed, press-disturbed, and intermediately disturbed systems would favour C, S and R strategies, respectively. However, several limitations remain— including the use of synthetic wastewater (excluding immigration effects), reliance on read-based metagenomics (which limits taxonomic resolution and functional linkage), small reactor volumes (15 mL), unequal numbers of disturbance events across experimental groups, and a labour-intensive analytical pipeline. We have since introduced a new computational tool, MicroEcoTools (Neshat, Santillan and Wuertz, 2025), to apply the CSR framework to disturbance experiment datasets in a reproducible fashion.

The aim of this work was to test the application of the CSR life-history strategy framework in anaerobic digestion bioreactors (also known as anaerobic digesters) fed with waste activated sludge—thus incorporating immigration effects—and exposed to varied frequencies of disturbance while maintaining an equal number of disturbance events across treatment groups. To this end, replicate laboratory-scale (mesocosm scale) anaerobic digesters were operated for a duration of 90 days. The digesters were subjected to varied frequencies of disturbance (via biomass removal) at a constant incident number. The microbial community composition and functional traits were analysed using a genome-resolved metagenomics approach. In addition to assessing community composition and trait distributions, we evaluated community-level functions to identify potential trade-offs induced by disturbance.

## 2. Material and methods

### 2.1. Experimental design

A set of twelve laboratory-scale anaerobic digesters with a working volume of 4 L was inoculated using anaerobic sludge obtained from a full-scale anaerobic digester operated at 35°C fed with a 1:1 mixture of primary and waste activated sludge (PS and WAS). After inoculation, the solids retention time (SRT) was reduced from 15 - 25 to 10 days. To acclimate the sludge to the new conditions, digesters were operated for a period of 7 months prior to the experiment. After acclimation, sludge was extracted from all digesters, mixed and redistributed among digesters with minimum air exposure. After that, a second month-long acclimation was performed to ensure the operational stability of digesters.

The acclimated digesters were then assigned to four experimental groups, each undergoing five disturbance events at varied frequencies (Figure S1). Throughout the experiment, daily sampling was done to assess the performance of the digesters. Samples were analysed on the same day for COD (soluble and total), solids (total and volatile), cations and anions and biogas composition. To assess the community structure and perform functional analysis of the metagenomes, one sample per digester was collected 10 days after the last disturbance event. For community analysis, samples were removed using syringes to avoid air exposure and transferred into cryo-vials containing ZymoResearch shield buffer (ZymoResearch, CA, USA) to inactivate the microorganisms and prevent degradation of the nucleic acids. These samples were stored at -80 °C until nucleic acid extraction.

### 2.2. Nucleic acid extraction and whole-genome shotgun sequencing

For nucleic acid extraction, 0.5 mL of sludge was thawed and transferred into ZymoResearch bashing beads tubes (ZymoResearch, CA, USA) and subjected to 4×40 s at a speed of 6 m/s of beating using a FastPrep-24 5G MP Biomedicals bead beating machine (MP Biomedicals, Singapore). Samples were cooled on ice for 5 min after every beating cycle. After bead beating, samples were centrifuged at maximum speed. Then, 0.2 mL of the supernatant was used to purify DNA using the ZymoResearch MagBead DNA/RNA extraction kit (ZymoResearch, CA, USA). An additional ethanol wash step was added to the manufacturer’s manual for improved contaminant removal. The nucleic acids were purified using an automated pipeline on a Tecan M200 Evo (Tecan, Männedorf, Switzerland) liquid handler. The quality of the extracted DNA was assessed using Nanodrop (Thermo Fisher Scientific, MA, USA), Qubit 4 (Thermo Fisher Scientific, MA, USA) and Agilent Tapestation (Agilent technologies, CA, USA) using a genomic DNA tape. Library preparation was performed using the Illumina TruSeq nano library preparation kit (Illumina, CA, USA). The libraries were sequenced on an Illumina Hiseq-X sequencing platform (Illumina, CA, USA) to obtain about 30 M paired-end reads per sample.

### 2.3. Genome-resolved bioinformatics and functional analysis

Changes in the microbial communities were assessed using a genome-resolved bioinformatics pipeline (Neshat *et al*., 2024). Briefly, raw reads were trimmed to remove sequencing adapters and low-quality reads using Trimmomatic v0.36 (Bolger, Lohse and Usadel, 2014) in paired-end mode (default parameters except for PE LEADING:3 TRAILING:3 SLIDINGWINDOW:4:15 MINLEN:36). Quality control for reads after and before trimming was performed using FastQC (Andrews, 2010). Trimmed reads from the same experimental group were then assembled into contigs using Spades v3.13 (Bankevich *et al*., 2012). After that, the raw reads were mapped onto the assembled contigs using BBMap (Bushnell, 2014). The mapping results were summarised using a Metabat2 function (Kang *et al*., 2019) (jgi_summarize_bam_contig_depths function). Metabat2 (Kang *et al*., 2019) was used to generate genome bins using coverage stats with default parameters. Generated metagenome-assembled genomes (MAGs) were dereplicated (at species level) and quality filtered using dRep v2.3.2 (Olm *et al*., 2017) (lineage workflow with -nc 0.5 -pa 0.9 -sa 0.99 -comp 50 -con 10). Taxonomy classification was performed using GTDB-tk v1.5 (Chaumeil *et al*., 2020). Relative abundances based on coverage were calculated using CoverM (Aroney *et al*., 2025) on genome mode using 20 M subsampled reads (mapping was performed using BBMap software package (Bushnell, 2014)). Functional annotation was performed using eggNOG-mapper v2.1.2 (Cantalapiedra *et al*., 2021) and the eggNOG 5.0 (Huerta-Cepas *et al*., 2019) database on the multi-FASTA files generated from the MAGs with at least MIMAG (Bowers *et al*., 2017) medium-quality. Seal function from BBMap (Bushnell, 2014) package was utilised to calculate the coverage of the annotated open reading frames (ORF) predicted by prodigal in eggNOG-mapper run (using 20M subsampled reads). Functional analysis results and coverage results were combined and summarised using an in-house script in the R environment. Briefly, annotated ORFs and mapping results were joined by ORF tags. After that, coverage was used as the weight variable to calculate the weighted counts for the cluster of orthologs genes (COG (Tatusov *et al*., 2003)) and carbohydrate-active enzyme (CAZy (Lombard *et al*., 2014)) categories, previously described as CATs (Santillan, Seshan, Constancias and Wuertz, 2019).

### 2.4. Physicochemical analysis

Ammonia and chemical oxygen demand analyses were performed using Hach kits (Hach COD HR and Hach TNT-plus ammonia HR kits [Hach, Loveland, CL, USA]) along with a Hach DR6000 spectrophotometer (Hach, Loveland, CL, USA). Solids (total, volatile and fixed) analysis was performed according to APHA AWA standard methods (Apha, 2005). Volumetric biogas production rate was measured using Bioprocess Instruments uFlow flowmeters equipped with 9 mL flow-cells (BP Instruments, Lund, Sweden). Analyses of biogas and volatile fatty acids composition were performed on a Shimadzu GC-2010 plus (Kyoto, Japan). A Shimadzu LC-20AD (Shimadzu, Kyoto, Japan) instrument was utilised for determination of anion and cation concentrations.

### 2.5. Statistical analysis

R v4.0.0 (Team, 2018) running on an Apple Mac OS Darwin 17.0 was utilised to perform all statistical analyses.

#### 2.5.1. Competitor – stress-tolerant – ruderal category assignment

MicroEcoTools (Neshat, Santillan and Wuertz, 2025), a computational tool for applying microbial ecology frameworks on computational data, was originally developed and utilised to classify MAGs and functional categories under the CSR framework. The CSR_assign function of this package assigns competitor, stress-tolerant and ruderal categories to organisms and functional traits abundant in the absence of, under intermediate and high levels of disturbance, respectively.

#### 2.5.2. Statistical comparison tests

For uni-variate comparison, the Welch-ANOVA (Welch, 1951) test employing the Benjamini-Hochberg method (Benjamini and Hochberg, 1995) for p-value adjustment was performed through the MicroEcoTools R package. In cases where the variance in at least one group was zero, ANOVA was employed to test the significance.

#### 2.5.3. Correlation analysis

Correlation analysis (Spearman’s rank) on community (coverage), functional (coverage weighted counts) and physicochemical data was performed using the Hmisc v4.6.0 (Harrell Jr and Harrell Jr, 2019) R package. P-value correction for multiple comparisons was performed using the Benjamini-Hochberg method. Results of the correlation analysis were plotted for pairs with significant correlations (adjusted p-value < 0.05). Cluster analysis (hierarchical clustering) was performed using the hclust base R function.

#### 2.5.4. Heatmaps

For heatmaps showing the abundance of MAGs, the depth of the tiles represents the relative abundance of each MAG (y-axis) in each digester (x-axis). For heatmaps showing the abundances of COG or CAZy categories, the weighted count of each category in each digester is shown as tile depth. For these heatmaps, x- and y-axes show frequency of disturbance and functional categories, respectively. To highlight the differences in each functional category (in COG and CAZy heatmaps), data were scaled per category. For all heatmaps, hierarchical double clustering was applied. Heatmaps were plotted using the pretty heatmaps (pheatmap) (Kolde and Kolde, 2015) R package.

#### 2.5.5. Multivariate analysis

Permutational multivariate analysis of variances (PERMANOVA) was employed to compare the community and functional compositions of the digesters across experimental groups. For this analysis, the adonis function of vegan v2.5.7 (Jari Oksanen, F. Guillaume Blanchet, Michael Friendly, Roeland Kindt, Dan McGlinn, Peter R. Minchin, R. B. O’Hara, Gavin L. Simpson, Peter Solymos and H. Stevens, 2019) R package using Bray-Curtis dissimilarity matrix was used with 999 permutations. The betadisper function in the same R package was used to assess the homogeneity of dispersion[39]. Constrained canonical analysis of principal coordinates (CAP) was performed to compare the communities under different disturbance levels. Physicochemical data were fitted to the CAP results and plotted as a biplot (only significant correlations are plotted).

#### 2.5.6. Network analysis

To assess the relationships between the community members and functional categories, correlation network analysis was performed. Network graph construction and visualisation was done using igraph v1.2.11 (Csardi G, 2006) and visNetwork v2.1.0 (Almende, Thieurmel and Robert, 2019) R packages.

## 3. Results

### 3.1. Disturbance frequency modulated the community-level functions

Community-level functional traits were affected differently by the disturbance depending on its frequency (Table 1). The highest biogas yield, solids content (volatile and total), total alkalinity and sulfate concentration values were observed in undisturbed reactors. On the other extreme end of the disturbance range, the highly disturbed reactors displayed the highest percentage of volatile matter removal (VMR). These reactors showed the lowest solids content, ammonia and total alkalinity values. The communities disturbed at intermediate frequencies of disturbance showed variable performance. While solids content values in those reactors were comparable, biogas production yield, VMR and ammonia concentration fluctuated. Notably, VMR was significantly higher in reactors exposed to a low frequency of disturbance compared to undisturbed reactors. Furthermore, ammonia and sulfate concentrations were the lowest in reactors disturbed at a low frequency of disturbance (2 disturbed days during 10 days of operation). The differences in community-level functions across experimental groups were statistically significant except for volatile fatty acids concentration (acetate, butyrate, propionate and valerate), pH, biogas volume and biogas composition (Table S1). Despite significant differences between alkalinity values across the experimental groups, the pH values were similar. All the reactors actively produced biogas and degraded volatile matter.

**TABLE 1.**
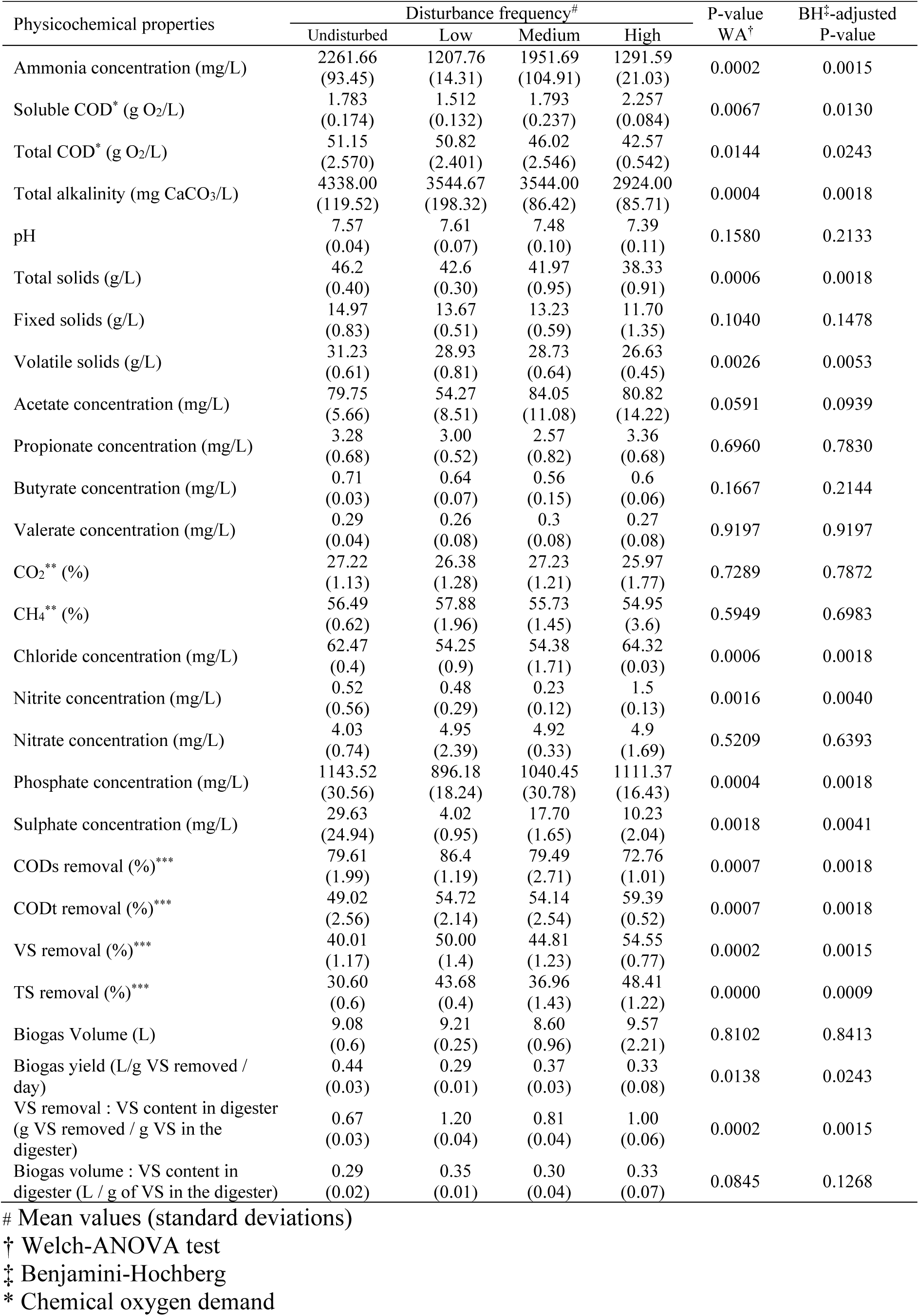
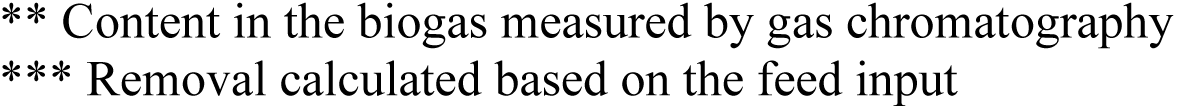
Summary of ecosystem traits and functions (process performance) and statistical comparison across digesters receiving different frequencies of disturbance after five disturbance events.

| Physicochemical properties | Disturbance frequency<sup>#</sup> |  |  |  | P-value<br>WA<sup>†</sup> | BH<sup>‡</sup>-adjusted<br>P-value |
| --- | --- | --- | --- | --- | --- | --- |
|  | Undisturbed | Low | Medium | High |  |  |
| Ammonia concentration (mg/L) | 2261.66<br>(93.45) | 1207.76<br>(14.31) | 1951.69<br>(104.91) | 1291.59<br>(21.03) | 0.0002 | 0.0015 |
| Soluble COD* (g O<sub>2</sub>/L) | 1.783<br>(0.174) | 1.512<br>(0.132) | 1.793<br>(0.237) | 2.257<br>(0.084) | 0.0067 | 0.0130 |
| Total COD* (g O<sub>2</sub>/L) | 51.15<br>(2.570) | 50.82<br>(2.401) | 46.02<br>(2.546) | 42.57<br>(0.542) | 0.0144 | 0.0243 |
| Total alkalinity (mg CaCO<sub>3</sub>/L) | 4338.00<br>(119.52) | 3544.67<br>(198.32) | 3544.00<br>(86.42) | 2924.00<br>(85.71) | 0.0004 | 0.0018 |
| pH | 7.57<br>(0.04) | 7.61<br>(0.07) | 7.48<br>(0.10) | 7.39<br>(0.11) | 0.1580 | 0.2133 |
| Total solids (g/L) | 46.2<br>(0.40) | 42.6<br>(0.30) | 41.97<br>(0.95) | 38.33<br>(0.91) | 0.0006 | 0.0018 |
| Fixed solids (g/L) | 14.97<br>(0.83) | 13.67<br>(0.51) | 13.23<br>(0.59) | 11.70<br>(1.35) | 0.1040 | 0.1478 |
| Volatile solids (g/L) | 31.23<br>(0.61) | 28.93<br>(0.81) | 28.73<br>(0.64) | 26.63<br>(0.45) | 0.0026 | 0.0053 |
| Acetate concentration (mg/L) | 79.75<br>(5.66) | 54.27<br>(8.51) | 84.05<br>(11.08) | 80.82<br>(14.22) | 0.0591 | 0.0939 |
| Propionate concentration (mg/L) | 3.28<br>(0.68) | 3.00<br>(0.52) | 2.57<br>(0.82) | 3.36<br>(0.68) | 0.6960 | 0.7830 |
| Butyrate concentration (mg/L) | 0.71<br>(0.03) | 0.64<br>(0.07) | 0.56<br>(0.15) | 0.6<br>(0.06) | 0.1667 | 0.2144 |
| Valerate concentration (mg/L) | 0.29<br>(0.04) | 0.26<br>(0.08) | 0.3<br>(0.08) | 0.27<br>(0.08) | 0.9197 | 0.9197 |
| CO<sub>2</sub>** (%) | 27.22<br>(1.13) | 26.38<br>(1.28) | 27.23<br>(1.21) | 25.97<br>(1.77) | 0.7289 | 0.7872 |
| CH<sub>4</sub>** (%) | 56.49<br>(0.62) | 57.88<br>(1.96) | 55.73<br>(1.45) | 54.95<br>(3.6) | 0.5949 | 0.6983 |
| Chloride concentration (mg/L) | 62.47<br>(0.4) | 54.25<br>(0.9) | 54.38<br>(1.71) | 64.32<br>(0.03) | 0.0006 | 0.0018 |
| Nitrite concentration (mg/L) | 0.52<br>(0.56) | 0.48<br>(0.29) | 0.23<br>(0.12) | 1.5<br>(0.13) | 0.0016 | 0.0040 |
| Nitrate concentration (mg/L) | 4.03<br>(0.74) | 4.95<br>(2.39) | 4.92<br>(0.33) | 4.9<br>(1.69) | 0.5209 | 0.6393 |
| Phosphate concentration (mg/L) | 1143.52<br>(30.56) | 896.18<br>(18.24) | 1040.45<br>(30.78) | 1111.37<br>(16.43) | 0.0004 | 0.0018 |
| Sulphate concentration (mg/L) | 29.63<br>(24.94) | 4.02<br>(0.95) | 17.70<br>(1.65) | 10.23<br>(2.04) | 0.0018 | 0.0041 |
| CODs removal (%)*** | 79.61<br>(1.99) | 86.4<br>(1.19) | 79.49<br>(2.71) | 72.76<br>(1.01) | 0.0007 | 0.0018 |
| CODt removal (%)*** | 49.02<br>(2.56) | 54.72<br>(2.14) | 54.14<br>(2.54) | 59.39<br>(0.52) | 0.0007 | 0.0018 |
| VS removal (%)*** | 40.01<br>(1.17) | 50.00<br>(1.4) | 44.81<br>(1.23) | 54.55<br>(0.77) | 0.0002 | 0.0015 |
| TS removal (%)*** | 30.60<br>(0.6) | 43.68<br>(0.4) | 36.96<br>(1.43) | 48.41<br>(1.22) | 0.0000 | 0.0009 |
| Biogas Volume (L) | 9.08<br>(0.6) | 9.21<br>(0.25) | 8.60<br>(0.96) | 9.57<br>(2.21) | 0.8102 | 0.8413 |
| Biogas yield (L/g VS removed / day) | 0.44<br>(0.03) | 0.29<br>(0.01) | 0.37<br>(0.03) | 0.33<br>(0.08) | 0.0138 | 0.0243 |
| VS removal : VS content in digester (g VS removed / g VS in the digester) | 0.67<br>(0.03) | 1.20<br>(0.04) | 0.81<br>(0.04) | 1.00<br>(0.06) | 0.0002 | 0.0015 |
| Biogas volume : VS content in digester (L / g of VS in the digester) | 0.29<br>(0.02) | 0.35<br>(0.01) | 0.30<br>(0.04) | 0.33<br>(0.07) | 0.0845 | 0.1268 |

<sup>#</sup> Mean values (standard deviations)
<sup>†</sup> Welch-ANOVA test
<sup>‡</sup> Benjamini-Hochberg
\* Chemical oxygen demand
\*\* Content in the biogas measured by gas chromatography \*\*\* Removal calculated based on the feed input

### 3.2. Three-way life-history strategies suggested by community structure and functional traits

The community structure reflects distinct patterns associated with different disturbance frequencies, as shown in the ordination plot from Canonical Analysis of Principal Coordinates (CAP) (Figure 1). Communities exposed to intermediate disturbance levels form a cohesive cluster, clearly separated from those under undisturbed or press disturbance conditions. These three clusters create a triangular arrangement, highlighting the impact of disturbance frequency on community structure. Separation along the primary axis (CAP1) differentiates intermediately disturbed communities from press-disturbed ones, while the secondary axis (CAP2) sets undisturbed communities apart from the rest.

**Figure 1.**
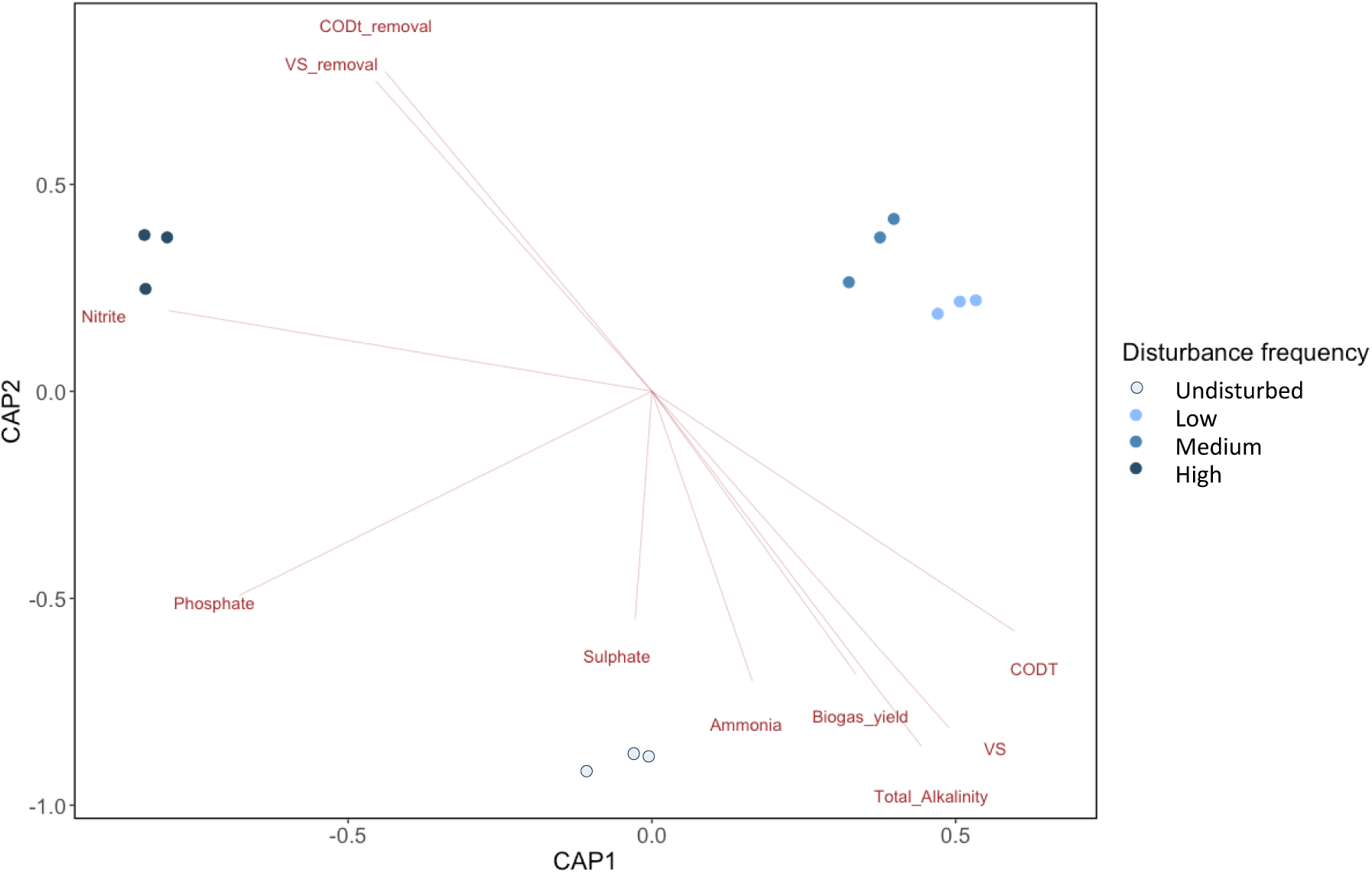
Canonical analysis of principal coordinates (CAP) on microbial community structure assessed using the genome-resolved metagenomics approach after five disturbance events at varied frequencies. The biplot vectors show the significant community-level trade-offs (Welch-ANOVA p-value < 0.05) in terms of total COD removal (CODt_removal [%]), volatile solids removal (VS_removal [%]), total COD content of the digesters (CODT), volatile solids content of the digesters (VS), total alkalinity value, biogas yield calculated as the volume of biogas produced per gram of volatile solids removed, ammonia, sulphate and nitrite concentrations.

Permutational analysis of variance on the communities also reveals that the communities are significantly different from each other using disturbance frequency as explanatory variable (PERMANOVA-F = 14.044, P-value = 0.001). Multivariate homogeneity of groups dispersion test (Anderson, 2006) was insignificant, suggesting the difference highlighted by PERMANOVA was not because of heteroscedasticity (PERMDISP F = 0.1814, P-value = 0.9061). The difference in community structure coincides with different community-level traits (vectors shown on the CAP plot), suggesting adaption of different CSR strategies. The correlation analysis on the function-community structure dataset shows significant correlations between disturbance level, functions and community structure supporting the above arguments (Figure S2).

Trade-offs were observed between solids content in the reactors (biomass production) and solids removal (consumption of organic materials for energy purposes). As highlighted by the correlation vectors, the disturbance frequency was positively correlated with the solids removal and negatively correlated with the solids content in the reactors. In addition, ion concentrations except for phosphate and nitrite were negatively correlated with the disturbance frequency.

### 3.3. Effect of disturbance frequency on community structure at genus level

The differences in community composition at the highest known taxonomy level across experimental groups is highlighted by a heatmap with double clustering (Figure 2 and figure S3 for top 25 and Figure S4 for top 100 taxa). The undisturbed conditions favoured taxa such as f_*Saprospiraceae*, g_*UBA8904* and g_*RZYY01*, while g_*Ottowia*, g_*MWFE01*, g_*Albidovulum*, f_*Rhodanobacteraceae*, g_*Sedimentibacter*, g_*Casimicrobium*, g_*OLB10*, g_*UBA1368* and g_*Sphaerochaeta* were more abundant in highly disturbed reactors. At low and medium disturbance frequencies, g_*Syntrophosphaera*, g_*Giesbergeria*, o_*UCB3* and g_*Phycicoccus-A* taxa were enriched. For the archaeal members of the community, g_*Methanosarcina* and *g*_*Methanothrix* were the dominant methanogens in all reactors (no difference across experimental groups), while g_*Methanospirillum* was only found in disturbed reactors. Changes in relative abundance of the top 25 genera across experimental groups are shown in Figure S3. The differences in the relative abundance of g_*Syntrophosphaera*, g_*DTU024*, g_*Lenti-01*, g_*Ottowia*, g_*DMER64*, f_*GCA-2699445*, g_*RZYY01*, g_*OLB10*, g_*Methanospirillum*, f_*UBA10799*, g_*Rectinema* and g_*Casimicrobium* taxa across experimental groups were significant based on Welch-ANOVA (Table S2). In addition, there were strongly significant correlations between the community members, resulting in three large clusters (Figure 3 and Figure S4. The community members (at genus level) were classified under the CSR framework (Table S2).

**Figure 2.**
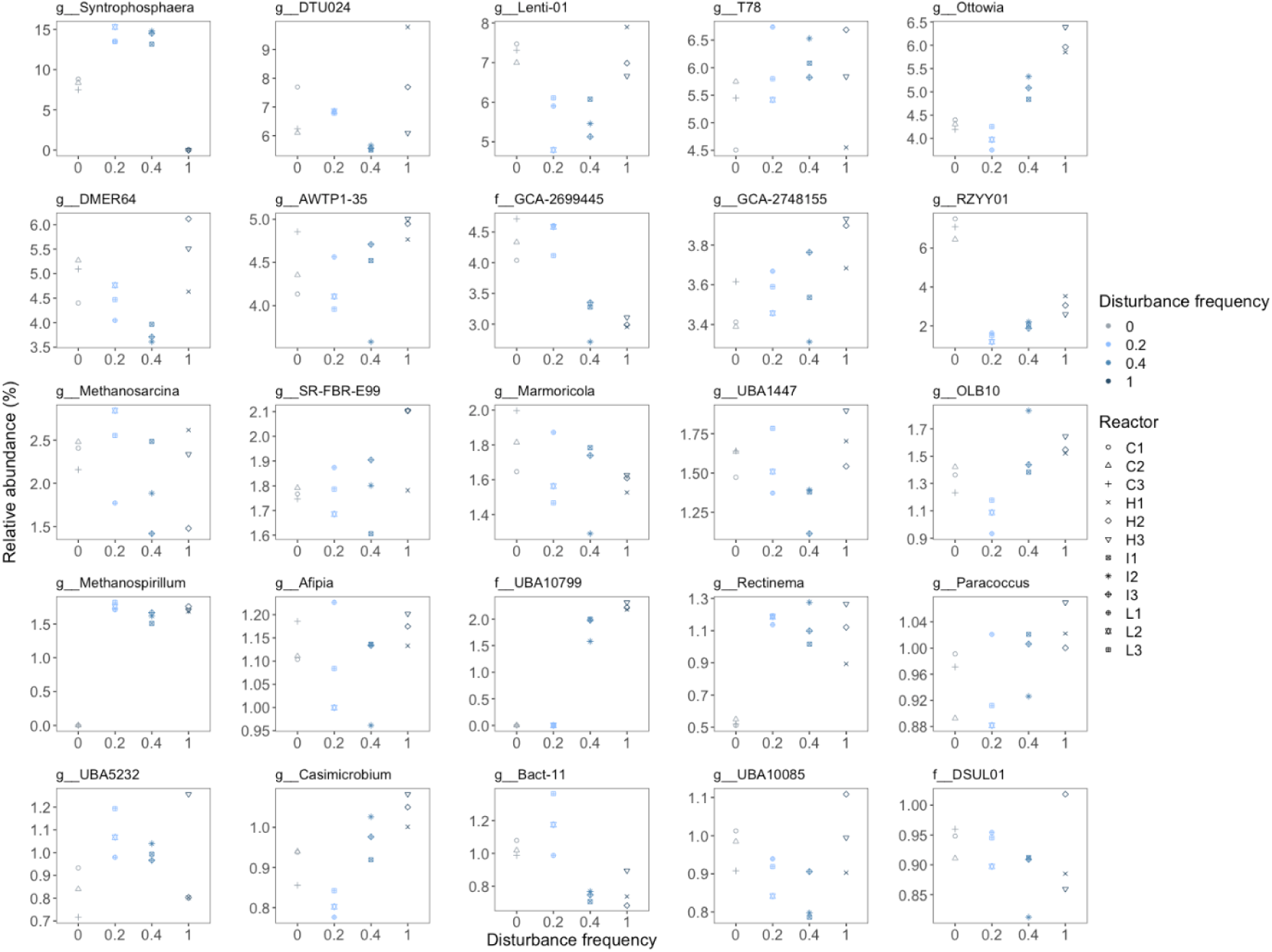
Relative abundance of the top 25 genera in replicate digesters (n=3) across experimental groups[18] based on the MAGs coverage.

**Figure 3.**
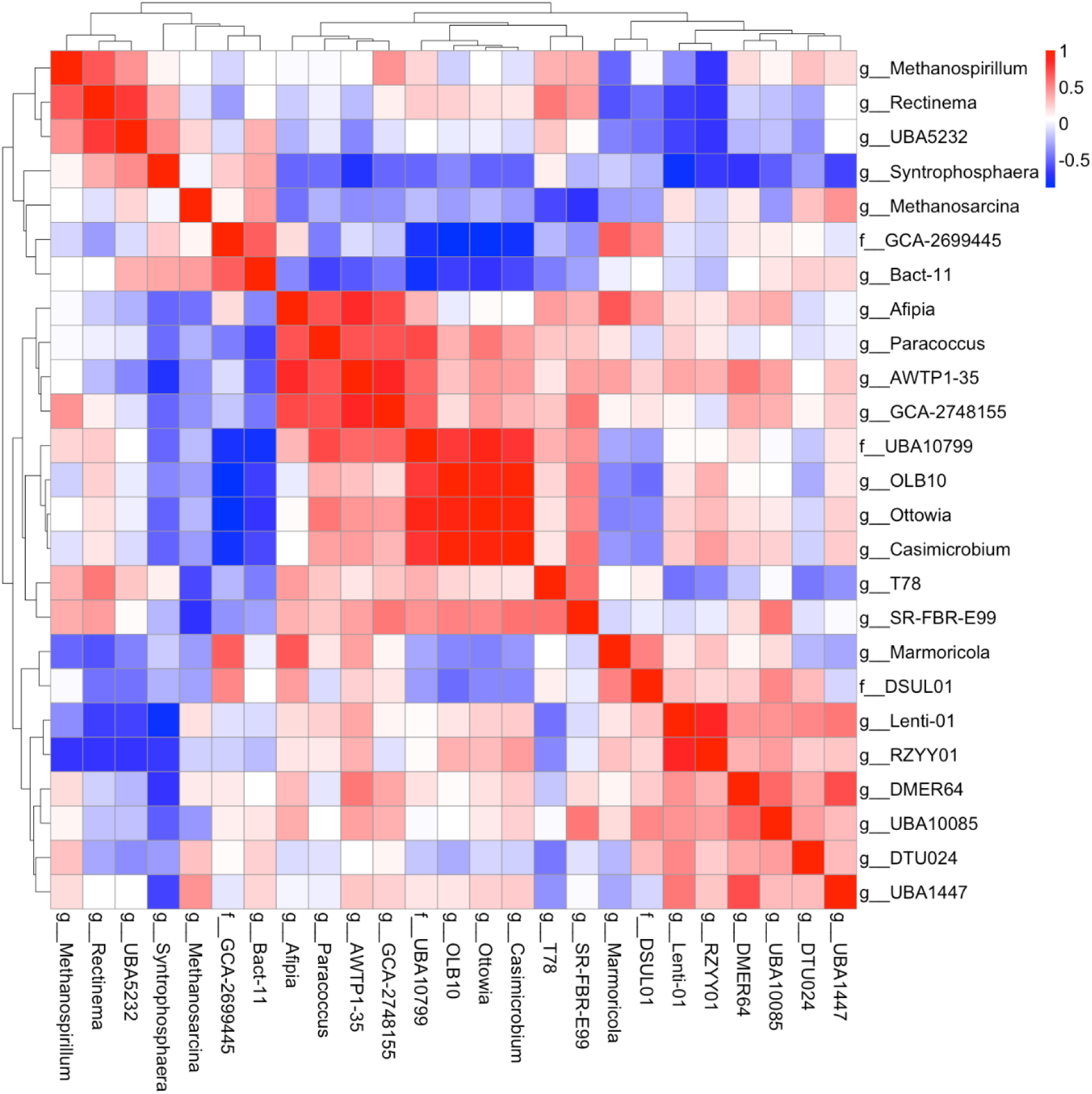
Spearman’s correlations for the 25 most abundant taxa (MAGs) with double clustering. Taxa are listed using their highest known taxonomy level (g: genus, f: family, o: order). The colour of the tiles shows the Spearman’s r value.

### 3.4. Functional potentials enriched by varying frequency of disturbance

Functional trait profiles diverged across disturbance regimes, with intermediately disturbed communities showing distinct patterns from undisturbed and press-disturbed groups. Cluster analysis of COG category abundances revealed a clear separation, with intermediate communities forming a cohesive group distinct from the other two (Figure 4, Figure S5). This pattern was supported by PERMANOVA (F = 52.04, P = 0.001), indicating a significant effect of disturbance frequency on functional composition, while PERMDISP confirmed homogeneity of group dispersions (F = 0.54, P = 0.67; Table S3, S4). Functional divergence was further reflected in the abundance of carbohydrate-active enzymes (CAZy), which also varied significantly among groups (Figure S6; Tables S3, S4). To explore life-history strategy patterns, COG and CAZy categories were classified under the CSR framework using the MicroEcoTools R package (Tables S5, S6).

**Figure 4.**
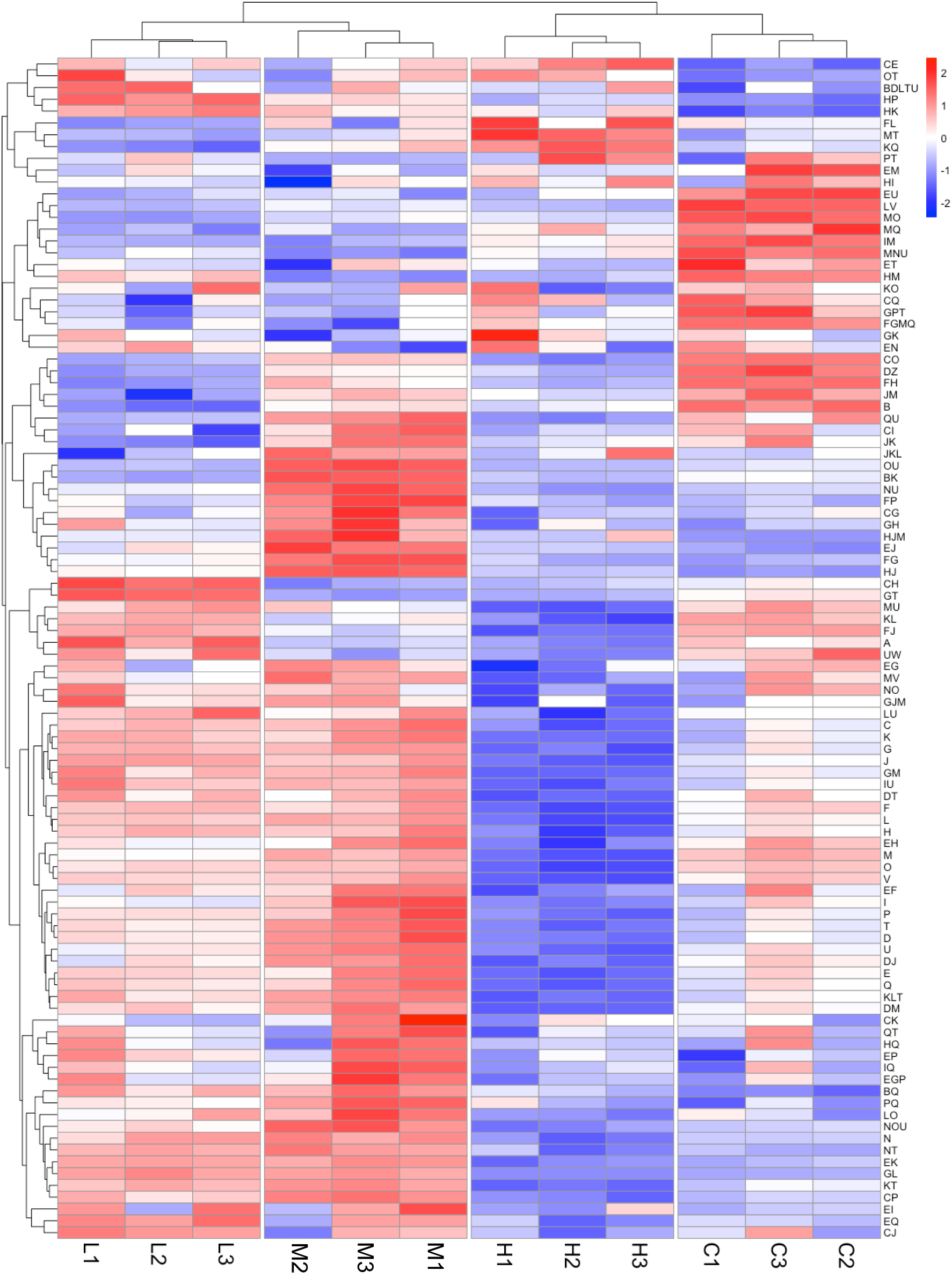
Abundance of the 100 most common COG categories annotated in the metagenomes across experimental groups. Only the genes present in the metagenomes with at least 10X coverage were included in this analysis. The x-axis shows the frequency of disturbance (C: undisturbed, L: low, M: Medium, H: High; numbers refer to the replicate digesters). The y-axis presents the COG categories (see Table S5 for COG category descriptions). The tile colours represent weighted counts that were Z-scale normalized across reactors for each COG category—meaning each count had its category’s average subtracted and was then divided by that category’s standard deviation.

### 3.5. Network analysis highlights a three-way life-history strategy in communities under varied frequencies of disturbance

Correlation network analysis revealed three distinct clusters for both the community structure and functional potentials annotated with the COG database (Figure 5). These clusters emerged when considering only strong, significant correlations (Spearman’s r > 0.5, p < 0.05), highlighting coherent groupings of taxa and functions shaped by disturbance frequency.

**Figure 5.**
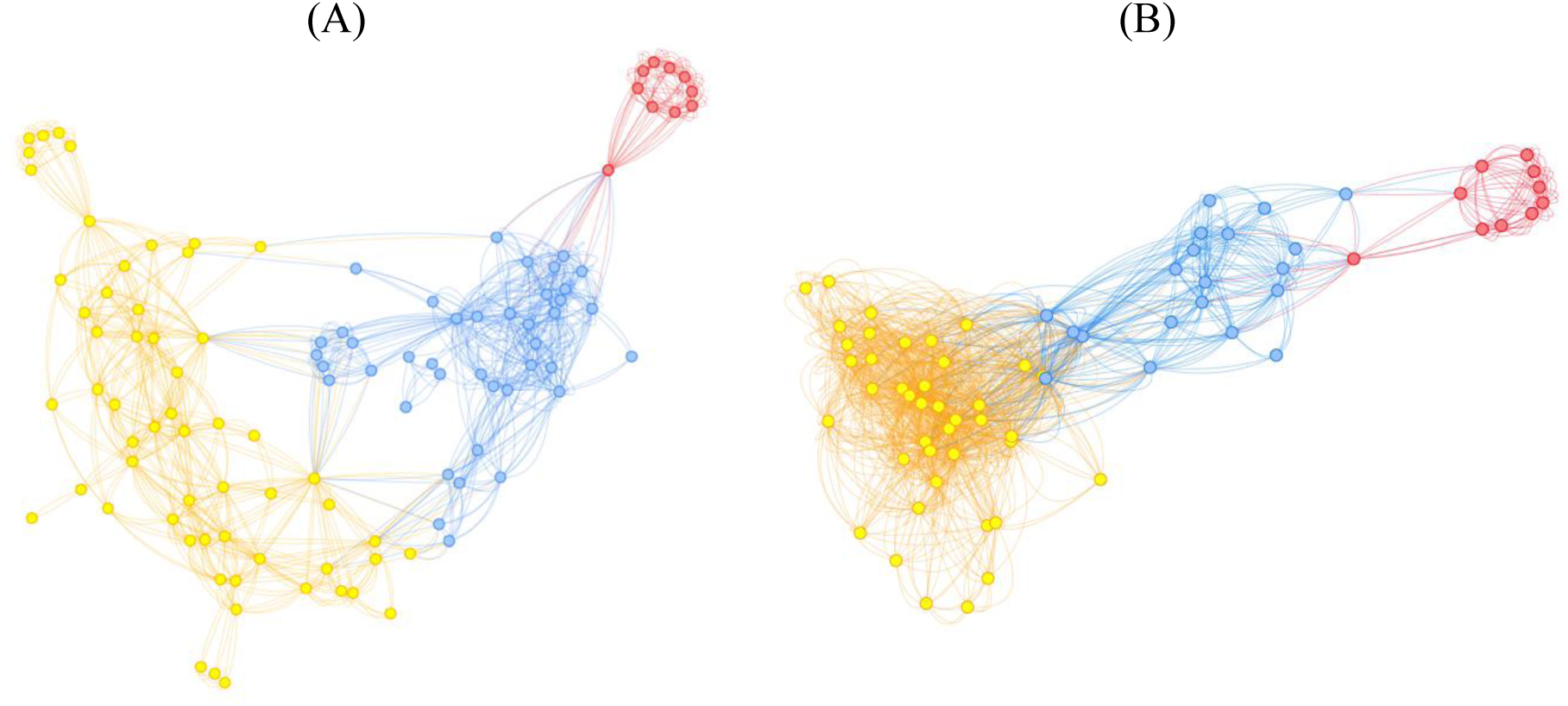
Correlation network analysis showing separate clusters using modularity of the network graph for taxa and functional potentials annotated with the COG database. (A) Community members at genus level; (B) COG functional categories with at least 10X coverage. The nodes in the network graphs show MAGs at the genus level or COG functional categories. Edges in the network graphs represent significant strong positive correlations (p-value < 0.05 and Spearman’s ρ > 0.5). Colours of the nodes and edges were determined with a modularity metric in the network graph using the resolution R package (Lambiotte, Delvenne and Barahona, 2008).

## 4. Discussion

### 4.1. Adoption of the three-way life-history strategies at community-level

Digesters that were disturbed at different frequencies at constant incidence separated in terms of community structure, and functional traits. Ordination analysis (CAP, Figure 1) revealed three distinct clusters aligned with the CSR life-history strategy triangle, indicating the emergence of competitor, ruderal, and stress-tolerant strategies. They also resulted in trade-offs in process performance, as seen in the CAP biplot. Together, these results support the adoption of three-way life-history strategies by the communities under disturbance, similar to the CSR framework (Grime, 1977). Ordination analyses have been previously employed to identify three-way life-history strategies in different environments, including activated sludge bacterial communities under disturbance (Santillan, Seshan, Constancias and Wuertz, 2019), fish communities (Pecuchet *et al*., 2017), coral reefs (Darling *et al*., 2012) and soil microbiome (Piton *et al*., 2023). To the best of our knowledge, the present study represents the first use of this framework in an open anaerobic digestion ecosystem using MAGs for assessing the community structure. The CSR classification encompassed 146 archaeal and bacterial genera, 25 ecosystem functions, and 201 COG trait complexes.

We propose that different frequency of disturbance, even at constant incidence, caused the adoption of different life-history strategies by the studied communities. The CSR strategies - competitor, ruderal and stress-tolerant - were adopted by undisturbed, intermediately disturbed and press disturbed communities, respectively. The community-level functions were distinct across categories: the highest biogas yield (the most efficient conversion of organics to biogas) was observed in competitors; soluble COD removal was highest in ruderals; and the VS content (equivalent to biomass) in reactors was higher in competitors and ruderals than in the communities categorised as stress-tolerant (highest frequency of disturbance). The adoption of three-way life-history strategies was also evident from the abundance of genes annotated with COG or CAZy categories in communities disturbed at different frequencies of disturbance. These patterns were supported by COG and CAZy gene abundances (Figures 4, S5, S6), network correlations (Figure 5), and Welch-ANOVA comparisons (Tables S5, S6), highlighting functional trade-offs aligned with life-history strategies.

### 4.2. Community structure, potential functions and community-level functional trade-offs unified under the CSR framework

Following the approach of Santillan et al. (Santillan, Seshan, Constancias and Wuertz, 2019), the community structure, potential functions, and community-level traits were combined into a conceptual CSR framework (Figure 6). While our study explicitly manipulated disturbance frequency, we recognize—consistent with Grime’s original formulation (Grime, 1977) —that CSR strategies arise from a combination of three ecological forces: disturbance, stress, and competition [12]. Different disturbance regimes can indirectly reflect these forces noted before (Santillan, Seshan, Constancias and Wuertz, 2019). For example, undisturbed reactors may reflect high competition due to resource accumulation and low diversity, while press-disturbed reactors may represent high-stress environments due to chronic biomass loss. Our schematic thus integrates this broader interpretation, providing a holistic representation of the ecological strategies adopted under varying conditions.

**Figure 6.**
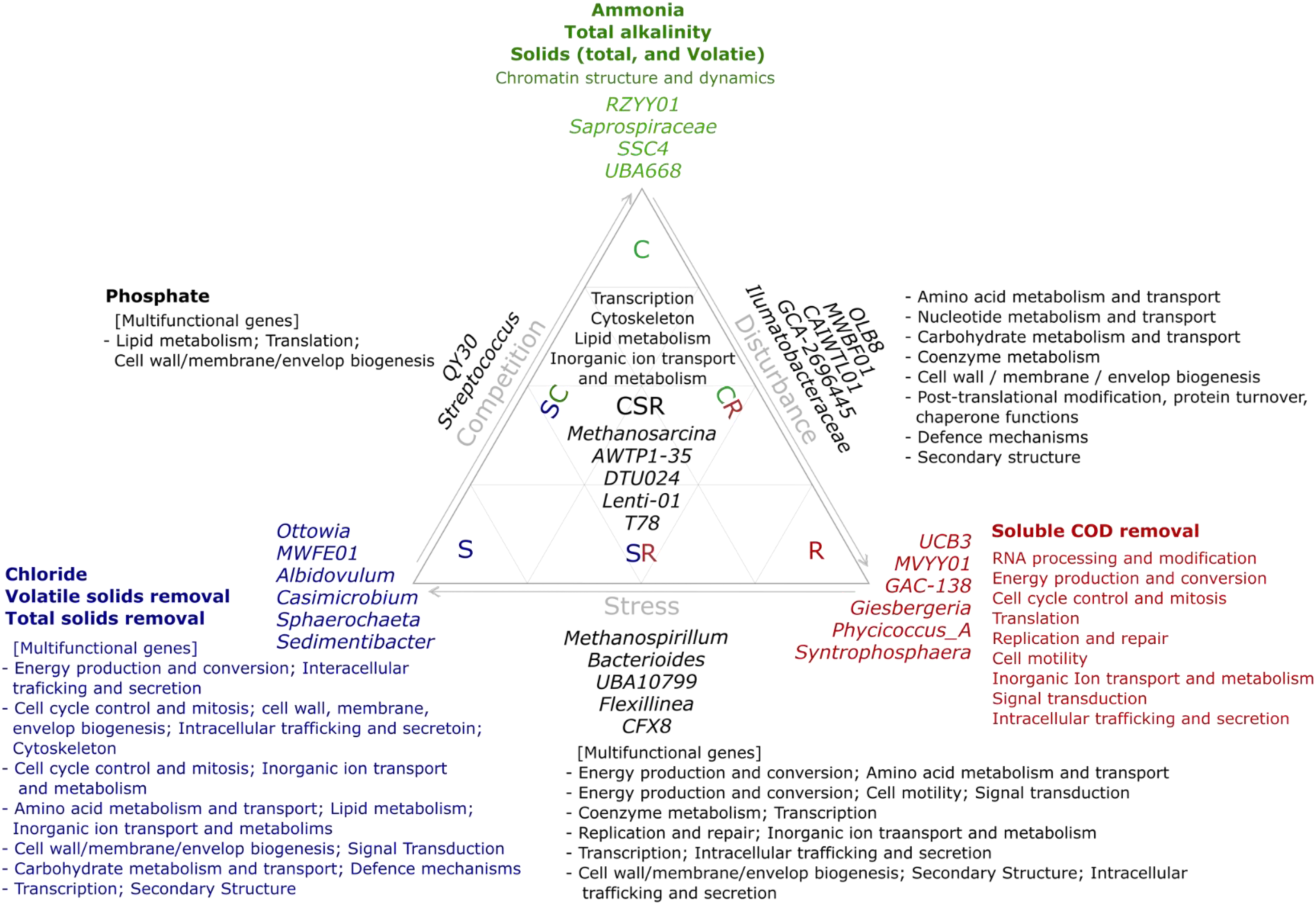
Distribution of taxa, COG functional categories and community-level functions in an anaerobic digestion ecosystem under varied frequencies of disturbance explained by the CSR framework. Traits classified under C, S and R categories are shown in green, blue and red colours, respectively. C: competitor; S: stress-tolerant; R: ruderal. Multi-functional genes are genes assigned to more than one COG category.

#### 4.2.1. Competitors (C category)

The high efficiency proposed by Grime and Pierce (Grime and Pierce, 2012) as one of the main indicators of competitive communities was observed in undisturbed communities in this study. Undisturbed reactors showed the highest biogas yield, suggesting better performance compared to disturbed reactors. In addition, the volatile solids content of the reactors (as a measure of biomass) was the highest in undisturbed reactors. This trait has also been proposed as one of the C-traits in activated sludge bacterial communities (Santillan, Seshan, Constancias and Wuertz, 2019) and in mycorrhizal fungal ecosystems (Chagnon *et al*., 2013). The genotypic functional analysis showed that genes assigned to more than one COG category (multi-functional genes) were dominant in undisturbed communities. These multi-functional genes are responsible for energy production and conversion, amino acid metabolism and transport, coenzyme metabolism, carbohydrate metabolism and transport, cell cycle and mitosis and cell wall/membrane/envelop biogenesis. Similar functional traits were reported before as C-traits (Santillan, Seshan, Constancias and Wuertz, 2019). Genes assigned to chromatin structure and dynamics were dominant in communities classified as competitor. This function has been reported to be related to growth rate (Travers and Muskhelishvili, 2005). Dominance of several taxa has been proposed as an indicator of competition (Santillan, Seshan, Constancias and Wuertz, 2019). Here, we observed that the undisturbed communities were dominated by several taxa—namely the genera *RZYY01, Saprospiraceae, SSC4 and UBA668*—whereas communities subjected to intermediate frequencies of disturbance were dominated by only one genus (*Syntrophosphaera*) with an average relative abundance of 14%.

#### 4.2.2. Ruderals (R category)

Ruderals are known for inefficiency in energy uptake (Grime and Pierce, 2012; Santillan, 2018), higher variability in ecosystem functions (Santillan, 2018; Santillan, Seshan, Constancias and Wuertz, 2019) and increased dissimilarities of communities across replicates exposed to the same frequency of disturbance (Santillan, 2018; Santillan, Seshan, Constancias and Wuertz, 2019). The reactors exposed to intermediate frequencies of disturbance in this study showed higher variability in terms of ecosystem functions (Table 1). In addition, the distances between replicate communities exposed to intermediate levels of disturbance in ordination analysis were greater than at the extreme ends of the disturbance range. Furthermore, it has been suggested that intermediate levels of disturbance create opportunities for less abundant community members, that is, seed-bank species, to outcompete dominant community members (Santillan, Seshan, Constancias and Wuertz, 2019; Santillan and Wuertz, 2022). The emergence of the genera including *Methanosprillium*, *Accumulibacter*, *GCA-2748155*, *Rectinema* and *MWFE01* was observed in disturbed reactors, while they were not abundant in undisturbed communities. Another reported trait for the ruderal category is a high biomass growth rate (Santillan, Seshan, Constancias and Wuertz, 2019). The VS content in intermediately disturbed reactors (disturbance frequencies of 0.2 and 0.4) was similar, despite having been disturbed at different frequencies. Considering the nature of the disturbance applied (biomass removal), biomass growth should have been promoted in reactors disturbed at a medium frequency for the VS content to be similar to the reactors disturbed at low disturbance frequency. This observation is consistent with Santillan et al. (Santillan, Seshan, Constancias and Wuertz, 2019) who reported higher biomass in the intermediately disturbed activated sludge communities compared to the undisturbed and press disturbed reactors. In a soil microbiome study (Fierer, Bradford and Jackson, 2007), high growth rate and yield were classified as copiotroph and oligotroph traits, respectively. Based on that study (Fierer, Bradford and Jackson, 2007), Krause et al. (Krause, 2014) categorised high growth rate as a competitor trait, which contrasts with the observation in this study.

Analysis of genotypic functional traits showed that genes assigned with only one COG category (specialised genes) were abundant in communities disturbed at intermediate frequencies (low and medium). This observation is in line with the notion that ruderals are inefficient (Krause, 2014; Santillan, 2018) as they have to express more genes to carry out a function. The same functions can be carried out by multi-functional genes in genera categorised as competitor. As such, energy production and conversion, translation, replication and repair, signal transduction, intracellular trafficking and secretion, cell cycle and control and cell motility were abundant COG categories in the communities labelled as ruderal. Cell motility function is another point of controversy trait-based studies. In accordance with the observation from this study, Krause et al. (Krause, 2014) categorised cell motility under the ruderal category, while Santillan et al. (Santillan, Seshan, Constancias and Wuertz, 2019) found that cell motility aggregated functions were abundant in stress-tolerant activated sludge communities. It was suggested that these traits are shared between S and R categories (Santillan, 2018; Santillan, Seshan, Constancias and Wuertz, 2019).

#### 4.2.3. Stress-tolerant (S category)

Organisms under high stress conditions are known to spend more energy on maintenance related functions to survive the harsh environmental conditions (Krause, 2014; Santillan, Seshan, Constancias and Wuertz, 2019). This key trait results in low biomass levels. The press disturbed communities in this study had the lowest biomass compared to others, which may have been due to the high stress condition or simply the result of the nature of the disturbance (biomass removal); hence we cannot conclude with certainty which factor was responsible for the low biomass in those communities. Remarkably, the genes assigned to multiple COG categories (multi-functional genes) were also more abundant in reactors exposed to a high frequency of disturbance. Multifunctional genes related to cell cycle control and mitosis, intracellular trafficking and secretion, cell wall/membrane/envelop biogenesis and inorganic ion transporters were the most abundant COG categories in the communities categorised as stress-tolerant. Similar traits (membrane chemistry and uptake system (Krause, 2014), transport and ATP binding cassette transporter genes (Santillan, Seshan, Constancias and Wuertz, 2019)) were classified as stress-tolerant in previous works.

#### 4.2.4. Inter-class categories (competitor–ruderal [CR], competitor–stress-tolerant [CS], ruderal-stress-tolerant [RS] and competitor-stress-tolerant-ruderal [CSR])

The majority of the microorganisms and genotypic traits in this study were classified into inter-categories —CR, CS, RS, or CSR—indicating either their abundance across more than one disturbance regime or a lack of significant difference among groups. Considering that the ecosystem was functional in all experimental groups, it can be assumed that the functions abundant in all categories (CSR) were essential for the core ecosystem. Diminishing those functions in an ecosystem like AD could have resulted in a breakdown. Intermediate classifications (e.g., CR, RS) reflect functional overlap or shared adaptive strategies between life-history categories. Inter-category traits have been reported in previous studies, for example, classification of cellular component organisation (Santillan, Seshan, Constancias and Wuertz, 2019) under the CR category, chemotaxis (Santillan, Seshan, Constancias and Wuertz, 2019) and sporulation (Santillan, Seshan, Constancias and Wuertz, 2019) related functions as SR and hyphal fusion (De La Providencia *et al*., 2005; Chagnon *et al*., 2013), growth rate (Hart and Reader, 2005; Chagnon *et al*., 2013) and cell division (Santillan, Seshan, Constancias and Wuertz, 2019) as CS. Additionally, some traits—such as cell motility (Krause, 2014; Santillan, Seshan, Constancias and Wuertz, 2019) and sporulation (Chagnon *et al*., 2013; Krause, 2014; Santillan, Seshan, Constancias and Wuertz, 2019)—have been classified under different categories in different studies. These discrepancies may have different roots; those traits arise because such traits are selected under multiple ecological pressures (for example, high disturbance and high stress), or due to methodological issues such as arbitrary cutoffs or insufficient statistical power. To overcome the classification errors, we employed the CSR_Simulation function of the MicroEcoTools package (Neshat, Santillan and Wuertz, 2025), which enables quantitative and reproducible assignment of traits to CSR categories through robust statistical comparisons across experimental groups.

### 4.3. Simulation-based classification further confirmed the community members, functions and traits assignments under the CSR framework

The misclassification of traits and genera across studies using different trait-based frameworks motivated us to develop a trait classification method based on pairwise comparison, considering the false discovery rate across the disturbance range, which was released as part of the MicroEcoTools R package (Neshat, Santillan and Wuertz, 2025). It was used to classify genera, functional traits and community-level functions in this study under the CSR framework (Tables S1, S2, S5 and, S6). Given the limited number of replicates in most studies, including the present one, one can increase the power of the classifications by employing simulation-based methods to test the probability of a trait being assigned to a CSR category. Hence, we applied another function of the MicroEcoTools, CSR_Simulation, that calculates the probability distribution of each trait/taxon having the observed abundance and uses that probability distribution to generate random communities. Then, simulated communities are used to calculate the probability of a trait/function. In most cases the analysis confirmed the classification assigned when only the observed data were considered but also suggested that traits/functions could have been classified under a different category based on additional experiments and minor variations in abundances. For example, the simulation showed that the genus *Albidovulum*, classified as R in this study, could be classified as CSR in 20% of the simulated communities. In another case, a genus in the family *Rhodanobacteraceae* was classified as CSR, but in 20% of the simulated communities it was classified as S. These differences were more obvious when categorising functional traits. For instance, RNA processing and modification was classified under the R category in the observed communities, but simulation showed that the probability of this category belonging to the R category was only 20%. These results further highlight the usefulness of *in silico* methods to address the limitations of a low number of experimental replicates.

## 5. Conclusions

This study demonstrates that life-history strategy frameworks, particularly Grime’s CSR model, provide a powerful conceptual tool for interpreting microbial community responses to disturbance in engineered ecosystems. By applying the CSR framework to anaerobic digestion—an ecologically and industrially important yet understudied ecosystem—we show that disturbance frequency at constant incident number can drive the emergence of distinct community strategies that align with competitive, ruderal, and stress-tolerant life-history patterns. Trait-based approaches, through the lens of CSR theory, allow for a more mechanistic understanding of community organization and functional trade-offs in response to disturbance. The identification of shared and intermediate strategies further reflects the complexity and adaptability of microbial life in dynamic environments. As the application of trait-based frameworks continues to expand in microbial systems, our study contributes to establishing their ecological validity in anaerobic systems and underscores their potential for guiding future research on microbial community resilience, succession, and ecosystem function in engineered settings.

## Data availability

DNA sequencing data are available in the NCBI BioProject database under accession number PRJNA723443.

## Acknowledgements

This research was supported by the Singapore National Research Foundation and Ministry of Education under the Research Centre of Excellence Program.

## Author Contributions

SN, ES and SW conceived the idea. SN designed and conducted the experiment and performed data processing and analysis. SN carried out the metagenomic bioinformatics. SW secured funding for the study. SN drafted the manuscript, and all authors contributed to its revision and editing.

