## Supplementary information for "Uncovering microbial life-history strategies under disturbance: a trait-based computational analysis of anaerobic systems"

### Supplementary figures

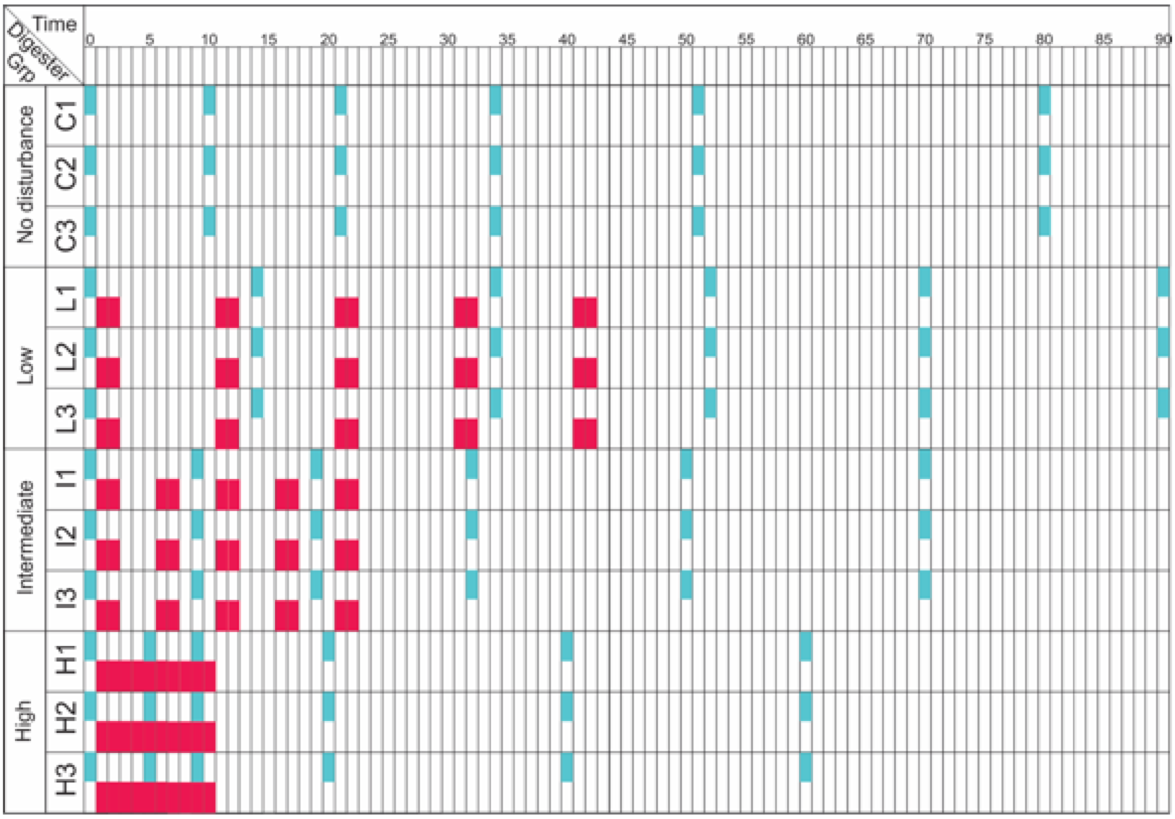


**Figure S1.** Experimental design and sampling scheme for the perturbation experiment. Disturbance events (reduction of solid retention time from 10 to 5 days equivalent to excess biomass removal) are shown in red colour and sampling time points for community analysis are shown in blue; the scale for time is days. Time points for community analysis were selected relative to the disturbance events; one time point before the start of the disturbances, one after the second disturbance event, one time point after the fourth disturbance event, one time point 10 days after the last disturbance event, and two time points 30, and 50 days after the last disturbance event.

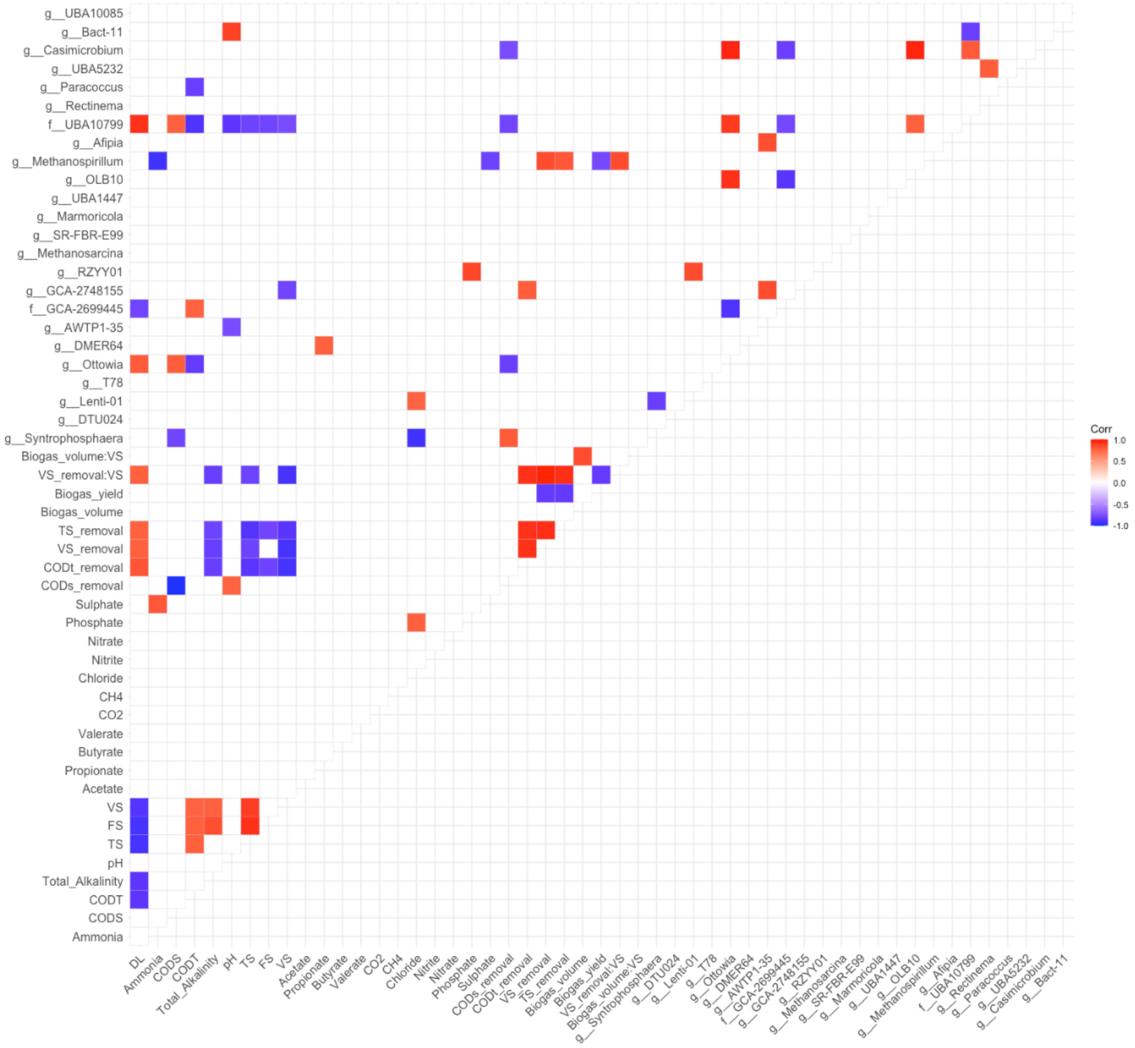


**Figure S2**. Correlogram of the community-level functions and the top-25 genera. Only strong significant correlations (P-value < 0.05 and Spearman's r > 0.6) were included. The colour of the tiles represents the spearman's r values.

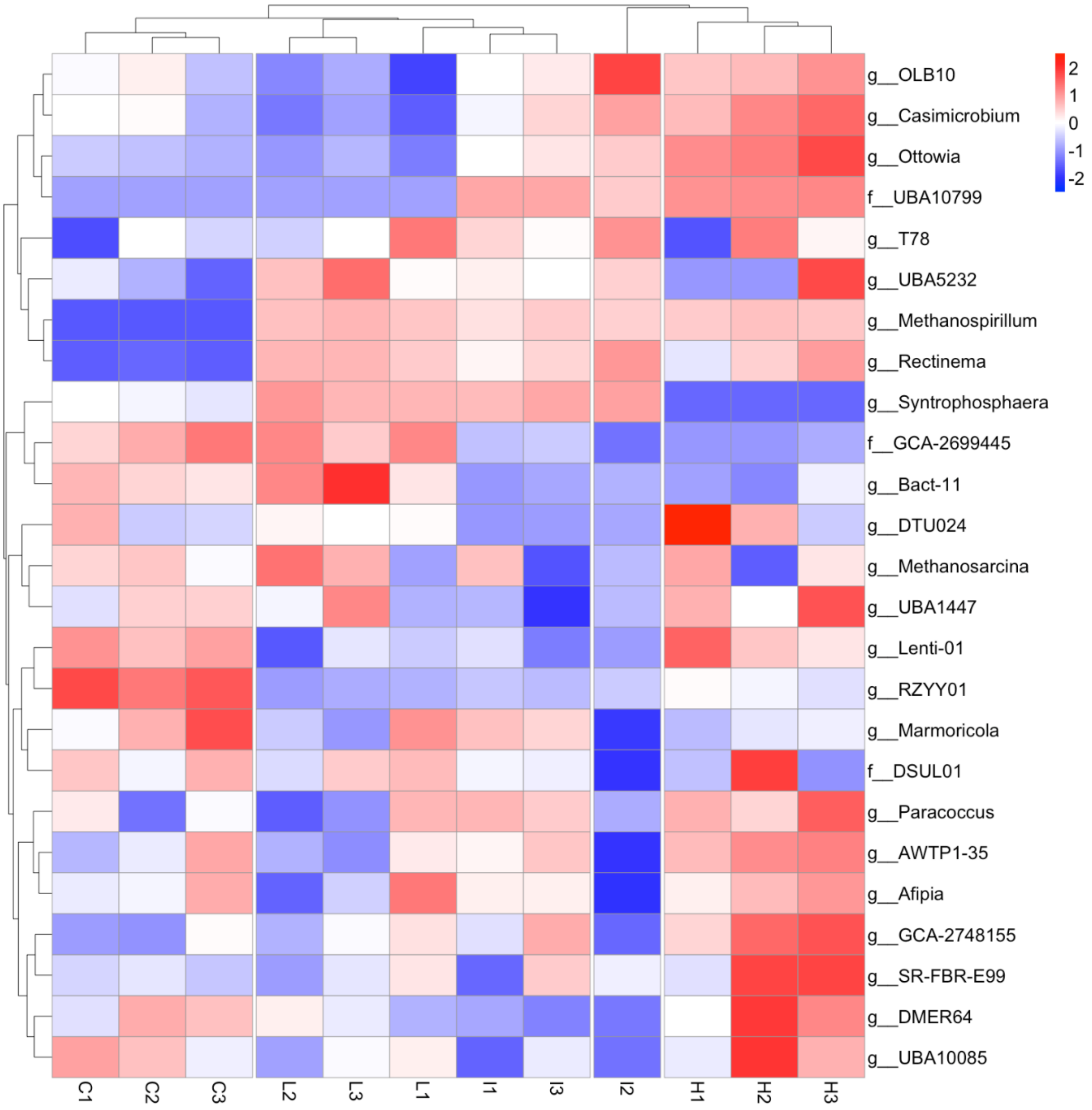


**Figure S3**. Clustered heatmap of the top 25 taxa (MAGs) analysed through the genome-resolved metagenomics approach across experimental groups. Communities disturbed at different frequencies are shown on x-axis (C: undisturbed, L: low [disturbance frequency = 0.2], I: intermediate [disturbance frequency = 0.4], press disturbed [disturbance frequency = 1]); numbers refer to the replicates; the y-axis shows the 25 most abundant taxa listed using the highest known taxonomy level in all samples (g: genus, f: family, o: order). Tile depth shows the Z-scale normalised coverage for each taxon in each reactor.

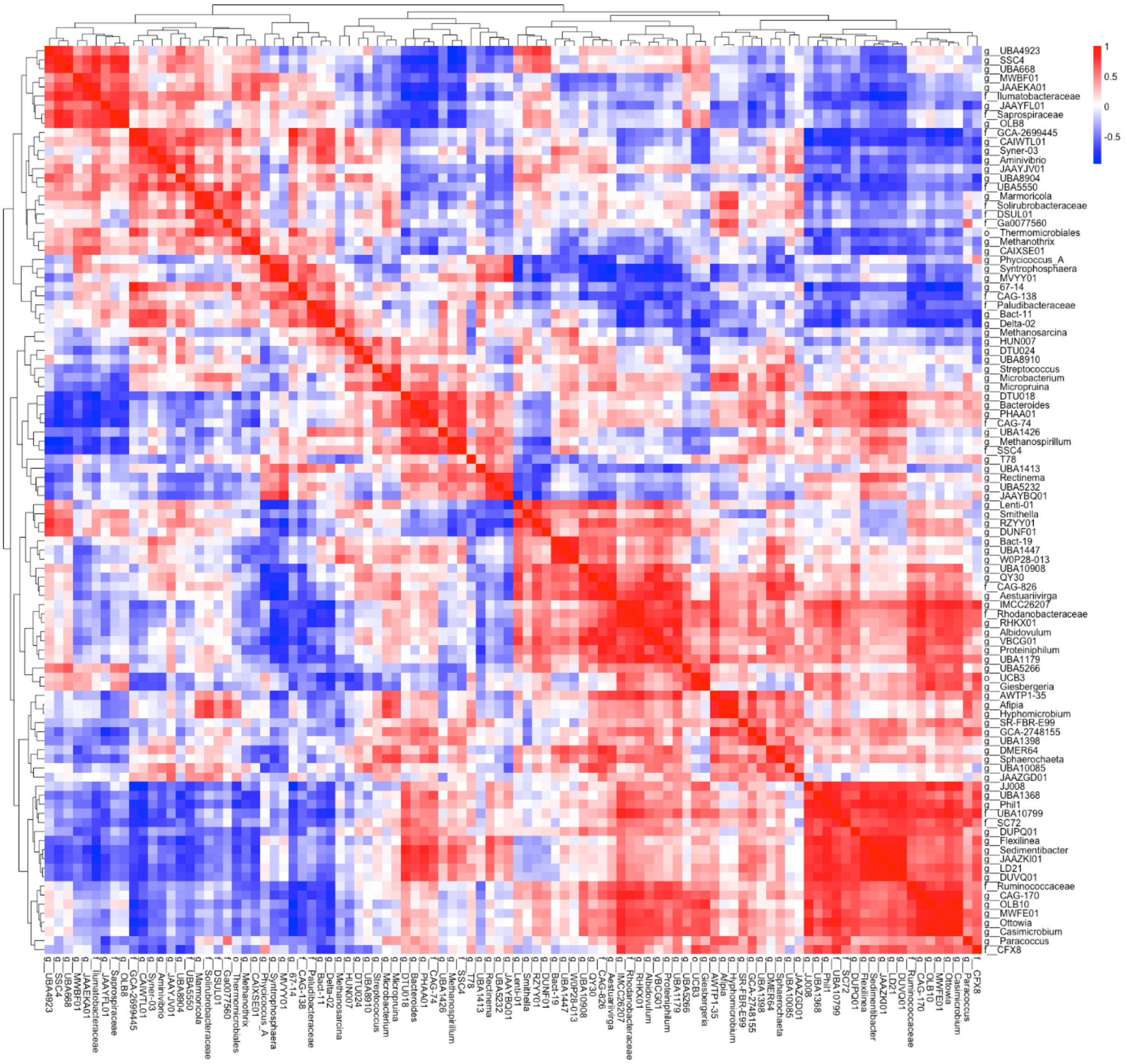


**Figure S4**. Correlation (Spearman's correlation) heatmap of top 100 genera with double clustering. Taxa are labelled using their highest known taxonomy level (g: genus, f: family, o: order).

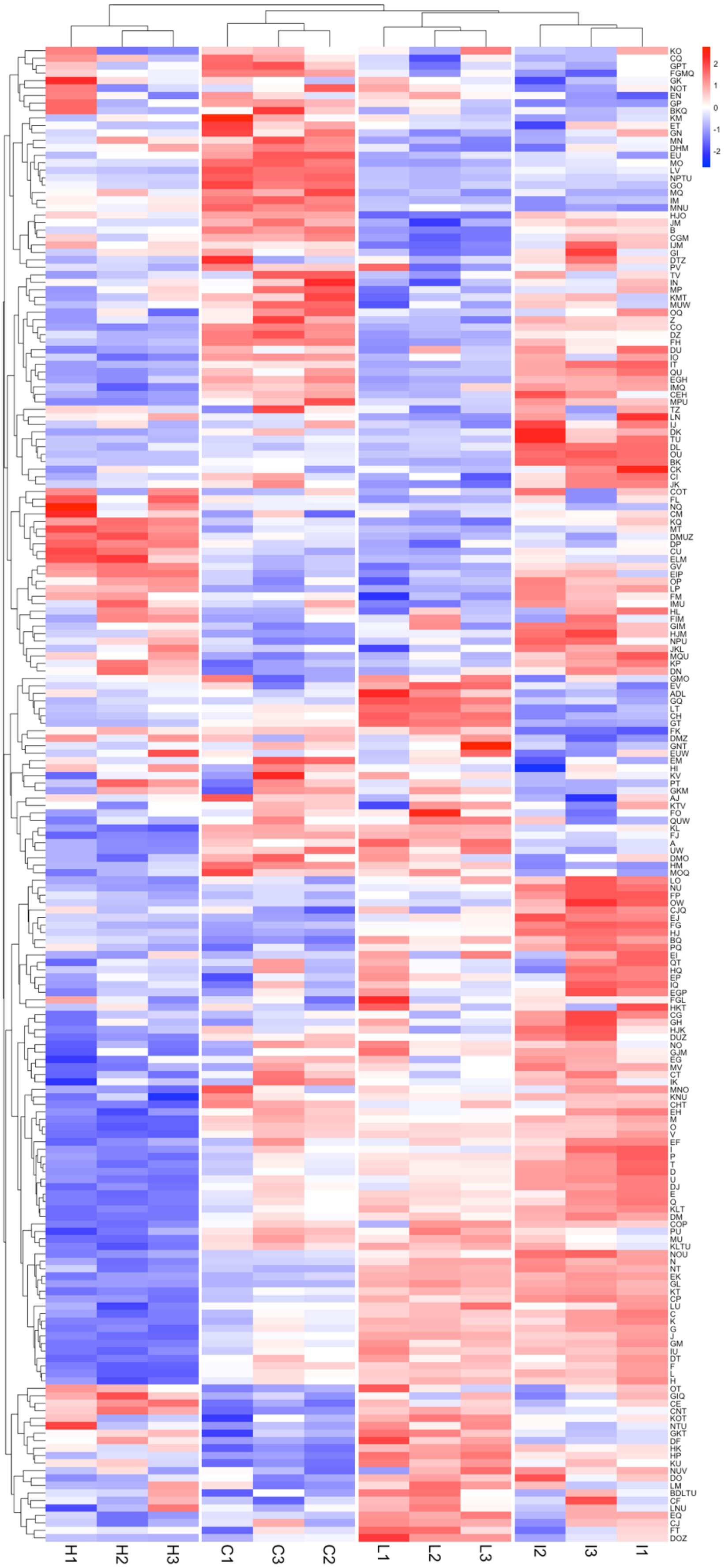
2
1
0
-1
-2
H1
H2
H3
C1
C3
C2
L1
L2
L3
I2
I3
I1

**Figure S5**. Abundance of the COG categories annotated in the metagenomes across experimental groups. The genes present in the metagenomes with at least 10X coverage were included in this analysis. The x-axis shows the metagenomes annotated in reactors disturbed at varied frequency of disturbance (C: undisturbed, L: low [disturbance frequency = 0.2], I: Intermediate [disturbance frequency = 0.4], press disturbed [disturbance frequency = 1]; numbers refer to the replicates). The y-axis presents the COG categories (see table S5 for COG category descriptions). Tile depth shows the Z-scale normalised (across reactors per COG category) weighted counts.

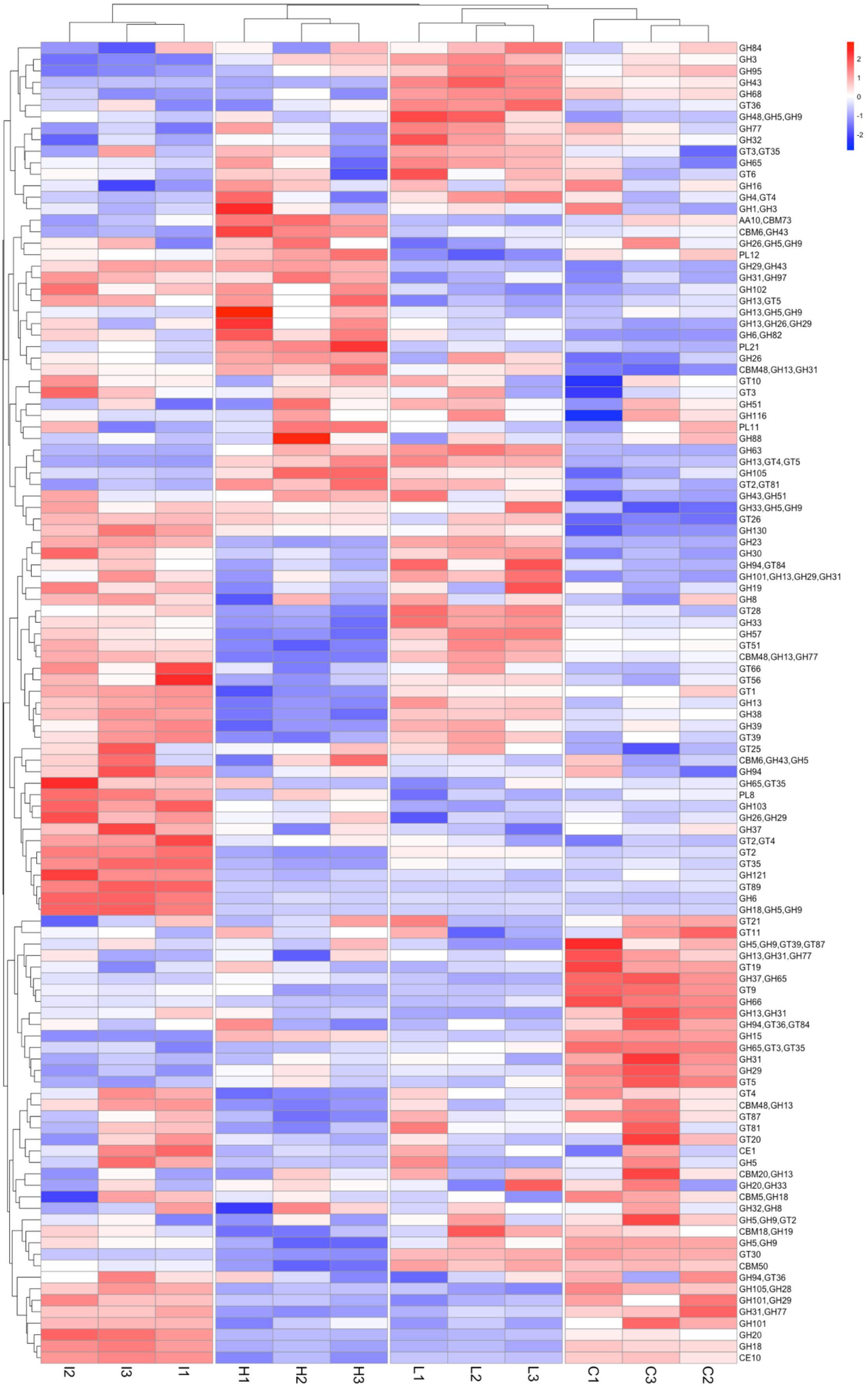


**Figure S6**. Abundance of the CAZy categories annotated in the metagenomes across experimental groups. The genes present in the metagenomes with at least 10X coverage were included in this analysis. The x-axis shows the metagenomes annotated in reactors disturbed at varied frequency of disturbance (C: undisturbed, L: low [disturbance frequency = 0.2], I: Intermediate [disturbance frequency = 0.4], press disturbed [disturbance frequency = 1]; numbers refer to the replicates). The y-axis presents the COG categories (see table 4.S6 for CAZy category descriptions). Tile depth shows the Z-scale normalised (normalisation was performed across reactors per COG category) weighted counts.

### Supplementary tables

**TABLE S1**. Summary result of statistical analysis for classification of the community-level traits under CSR framework

| Community trait | Welch ANOVA p-value | BH* adjusted p-value | Cr[§] | CR[§] | CS[§] | rR[§] | rS[§] | RS[§] | CSR[§] |
|---|---|---|---|---|---|---|---|---|---|
| Ammonia | 0.000 | 0.002 | C | C | C | R | S | R | C |
| Total alkalinity | 0.000 | 0.002 | C | C | C | N | R | R | C |
| TS | 0.001 | 0.002 | C | C | C | N | R | R | C |
| VS | 0.003 | 0.006 | C | C | C | N | R | R | C |
| Phosphate | 0.000 | 0.002 | C | C | N | R | S | S | CS |
| Nitrite | 0.002 | 0.004 | N | N | N | N | S | S | CSR |
| Sulphate | 0.002 | 0.004 | N | N | N | R | S | R | CSR |
| CODS | 0.007 | 0.014 | N | N | N | N | S | N | CSR |
| CODT | 0.014 | 0.024 | N | N | N | N | N | N | CSR |
| CODt removal | 0.014 | 0.024 | N | N | N | N | N | N | CSR |
| Biogas yield | 0.014 | 0.024 | N | N | N | N | N | N | CSR |
| Acetate | 0.059 | 0.094 | N | N | N | N | N | N | CSR |
| FS | 0.104 | 0.148 | N | N | N | N | N | N | CSR |
| pH | 0.158 | 0.213 | N | N | N | N | N | N | CSR |
| Butyrate | 0.167 | 0.214 | N | N | N | N | N | N | CSR |
| Nitrate | 0.521 | 0.639 | N | N | N | N | N | N | CSR |
| CH4 | 0.595 | 0.698 | N | N | N | N | N | N | CSR |
| Propionate | 0.696 | 0.783 | N | N | N | N | N | N | CSR |
| CO2 | 0.729 | 0.787 | N | N | N | N | N | N | CSR |
| Biogas volume | 0.810 | 0.841 | N | N | N | N | N | N | CSR |
| Valerate | 0.920 | 0.920 | N | N | N | N | N | N | CSR |
| CODs removal | 0.001 | 0.002 | R | N | C | r | R | R | R |
| TS removal | 0.000 | 0.001 | R | R | S | r | S | S | S |
| Chloride | 0.001 | 0.002 | C | C | S | N | S | S | S |
| VS removal | 0.000 | 0.002 | R | R | S | r | S | S | S |

§: Pairwise comparison between groups C: competitor (no disturbance); r ruderal (low frequency of disturbance); R ruderal (medium frequency of disturbance); S: stress-tolerant (high frequency of disturbance)
*: Benjamini-Hochberg

**TABLE S2**. Summary result of statistical analysis for classification of the community members (highest known taxonomy level) under CSR framework. Table generated using MicroEcoTools R package (Neshat, Santillan and Wuertz, 2025).

| TAXA | Welch ANOVA p-value | BH† adjusted p-value | CSR‡ | Probability calculated by bootstrapping (%) | | | | | | |
|---|---|---|---|---|---|---|---|---|---|---|
| | | | | C§ | R§ | S§ | CR§ | CS§ | SR§ | CSR§ |
| g_Syntrophosphaera | 0.000* | 0.000* | R | 0.00 | 100 | 0.00 | 0.00 | 0.00 | 0.00 | 0.00 |
| g_DTU024 | 0.000 | 0.001 | CSR | 0.00 | 0.00 | 0.00 | 0.00 | 0.00 | 0.00 | 100 |
| g_Lenti-01 | 0.024 | 0.034 | CSR | 0.00 | 0.00 | 0.00 | 0.00 | 0.00 | 0.00 | 100 |
| g_T78 | 0.418 | 0.449 | CSR | 0.00 | 0.00 | 0.00 | 0.00 | 0.00 | 0.00 | 100 |
| g_Ottowia | 0.003 | 0.006 | S | 0.00 | 0.00 | 100 | 0.00 | 0.00 | 0.00 | 0.00 |
| g_DMER64 | 0.037 | 0.052 | CSR | 0.00 | 0.00 | 0.00 | 0.00 | 0.00 | 0.00 | 100 |
| g_AWTP1-35 | 0.101 | 0.126 | CSR | 0.00 | 0.00 | 0.00 | 0.00 | 0.00 | 0.00 | 100 |
| f_GCA-2699445 | 0.006 | 0.010 | CR | 0.00 | 0.00 | 0.00 | 100 | 0.00 | 0.00 | 0.00 |
| g_GCA-2748155 | 0.130 | 0.156 | CSR | 0.00 | 0.00 | 0.00 | 0.00 | 0.00 | 0.00 | 100 |
| g_RZYY01 | 0.001 | 0.001 | C | 100 | 0.00 | 0.00 | 0.00 | 0.00 | 0.00 | 0.00 |
| g_Methanosarcina | 0.712 | 0.726 | CSR | 0.00 | 0.00 | 0.00 | 0.00 | 0.00 | 0.00 | 100 |
| g_SR-FBR-E99 | 0.462 | 0.485 | CSR | 0.00 | 0.00 | 0.00 | 0.00 | 0.00 | 0.00 | 100 |
| g_Marmoricola | 0.428 | 0.456 | CSR | 0.00 | 0.00 | 0.00 | 0.00 | 0.00 | 0.00 | 100 |
| g_UBA1447 | 0.171 | 0.198 | CSR | 0.00 | 0.00 | 0.00 | 0.00 | 0.00 | 0.00 | 100 |
| g_OLB10 | 0.019 | 0.029 | CSR | 0.00 | 0.00 | 0.00 | 0.00 | 0.00 | 0.00 | 100 |
| g_Methanospirillum | 0.000* | 0.000* | SR | 0.00 | 0.00 | 0.00 | 0.00 | 0.00 | 100 | 0.00 |
| g_Afipia | 0.526 | 0.548 | CSR | 0.00 | 0.00 | 0.00 | 0.00 | 0.00 | 0.00 | 100 |
| f_UBA10799 | 0.000* | 0.000* | SR | 0.00 | 0.00 | 1.20 | 0.00 | 0.00 | 98.80 | 0.00 |
| g_Rectinema | 0.000 | 0.000 | CSR | 0.00 | 0.00 | 0.00 | 0.00 | 0.00 | 0.00 | 100 |
| g_Paracoccus | 0.280 | 0.309 | CSR | 0.00 | 0.00 | 0.00 | 0.00 | 0.00 | 0.00 | 100 |
| g_UBA5232 | 0.238 | 0.268 | CSR | 0.00 | 0.00 | 0.00 | 0.00 | 0.00 | 0.00 | 100 |
| g_Casimicrobium | 0.008 | 0.013 | S | 0.00 | 0.00 | 75.22 | 0.00 | 0.00 | 0.00 | 24.78 |
| g_Bact-11 | 0.006 | 0.010 | CSR | 0.00 | 0.00 | 0.00 | 0.00 | 0.00 | 0.00 | 100 |
| g_UBA10085 | 0.177 | 0.204 | CSR | 0.00 | 0.00 | 0.00 | 0.00 | 0.00 | 0.00 | 100 |
| f_DSUL01 | 0.568 | 0.588 | CSR | 0.00 | 0.00 | 0.00 | 0.00 | 0.00 | 0.00 | 100 |
| g_Hyphomicrobium | 0.593 | 0.609 | CSR | 0.00 | 0.00 | 0.00 | 0.00 | 0.00 | 0.00 | 100 |
| f_Saprospiraceae | 0.034 | 0.048 | C | 94.91 | 0.00 | 0.00 | 0.00 | 0.00 | 0.00 | 5.09 |
| g_67-14 | 0.208 | 0.237 | CSR | 0.00 | 0.00 | 0.00 | 0.00 | 0.00 | 0.00 | 100 |
| o__UCB3 | 0.005 | 0.010 | R | 0.00 | 100 | 0.00 | 0.00 | 0.00 | 0.00 | 0.00 |
| g_Phycicoccus_A | 0.001 | 0.001 | R | 0.00 | 100 | 0.00 | 0.00 | 0.00 | 0.00 | 0.00 |
| g_Albidovulum | 0.005 | 0.009 | S | 0.00 | 0.00 | 79.62 | 0.00 | 0.00 | 0.00 | 20.38 |
| g_Proteiniphilum | 0.125 | 0.150 | CSR | 0.00 | 0.00 | 0.00 | 0.00 | 0.00 | 0.00 | 100 |
| g_MWFE01 | 0.002 | 0.004 | S | 0.00 | 0.00 | 100 | 0.00 | 0.00 | 0.00 | 0.00 |
| g_Giesbergeria | 0.002 | 0.004 | R | 0.00 | 100 | 0.00 | 0.00 | 0.00 | 0.00 | 0.00 |
| g_Syner-03 | 0.002 | 0.004 | CSR | 0.00 | 0.00 | 0.00 | 0.00 | 0.00 | 0.00 | 100 |
| f_Rhodanobacteraceae | 0.020 | 0.029 | CSR | 0.00 | 0.00 | 20.18 | 0.00 | 0.00 | 0.00 | 79.82 |
| g_JAAZGD01 | 0.935 | 0.942 | CSR | 0.00 | 0.00 | 0.00 | 0.00 | 0.00 | 0.00 | 100 |

| TAXA | Welch ANOVA p-value | BH† adjusted p-value | CSR‡ | Probability calculated by bootstrapping (%) | | | | | | |
|---|---|---|---|---|---|---|---|---|---|---|
| | | | | C§ | R§ | S§ | CR§ | CS§ | SR§ | CSR§ |
| f_Ga0077560 | 0.947 | 0.947 | CSR | 0.00 | 0.00 | 0.00 | 0.00 | 0.00 | 0.00 | 100 |
| g_Sedimentibacter | 0.000* | 0.000* | S | 0.00 | 0.00 | 100 | 0.00 | 0.00 | 0.00 | 0.00 |
| g_JAAYBQ01 | 0.009 | 0.015 | CSR | 0.00 | 0.00 | 0.00 | 0.00 | 0.00 | 0.00 | 100 |
| g_Methanothrix | 0.330 | 0.362 | CSR | 0.00 | 0.00 | 0.00 | 0.00 | 0.00 | 0.00 | 100 |
| g_LD21 | 0.037 | 0.052 | CSR | 0.00 | 0.00 | 0.00 | 0.00 | 0.00 | 0.00 | 100 |
| g_UBA1398 | 0.339 | 0.369 | CSR | 0.00 | 0.00 | 0.00 | 0.00 | 0.00 | 0.00 | 100 |
| g_UBA8904 | 0.068 | 0.087 | CSR | 0.00 | 0.00 | 0.00 | 0.00 | 0.00 | 0.00 | 100 |
| g_Smithella | 0.168 | 0.196 | CSR | 0.00 | 0.00 | 0.00 | 0.00 | 0.00 | 0.00 | 100 |
| g_Microbacterium | 0.441 | 0.466 | CSR | 0.00 | 0.00 | 0.00 | 0.00 | 0.00 | 0.00 | 100 |
| g_Sphaerochaeta | 0.012 | 0.019 | S | 0.00 | 0.00 | 100 | 0.00 | 0.00 | 0.00 | 0.00 |
| g_Flexilinea | 0.001 | 0.002 | SR | 0.00 | 0.00 | 0.00 | 0.00 | 0.00 | 100 | 0.00 |
| g_UBA1368 | 0.064 | 0.084 | CSR | 0.00 | 0.00 | 0.00 | 0.00 | 0.00 | 0.00 | 100 |
| g_Bact-19 | 0.002 | 0.004 | CSR | 0.00 | 0.00 | 0.00 | 0.00 | 0.00 | 0.00 | 100 |
| f_SC72 | 0.022 | 0.031 | S | 0.00 | 0.00 | 100 | 0.00 | 0.00 | 0.00 | 0.00 |
| g_Delta-02 | 0.009 | 0.014 | CSR | 0.00 | 1.10 | 0.00 | 0.00 | 0.00 | 0.00 | 98.90 |
| g_DTU018 | 0.048* | 0.065* | CSR | 0.00 | 0.00 | 0.00 | 0.00 | 0.00 | 0.00 | 100 |
| g_W0P28-013 | 0.105 | 0.129 | CSR | 0.00 | 0.00 | 0.00 | 0.00 | 0.00 | 0.00 | 100 |
| g_UBA1413 | 0.137 | 0.162 | CSR | 0.00 | 0.00 | 0.00 | 0.00 | 0.00 | 0.00 | 100 |
| g_PHAA01 | 0.007 | 0.012 | SR | 0.00 | 0.00 | 9.19 | 0.00 | 0.00 | 90.41 | 0.40 |
| g_Bacteroides | 0.011 | 0.017 | SR | 0.00 | 0.00 | 0.00 | 0.00 | 0.00 | 100 | 0.00 |
| f_Solirubrobacteraceae | 0.751 | 0.762 | CSR | 0.00 | 0.00 | 0.00 | 0.00 | 0.00 | 0.00 | 100 |
| g_QY30 | 0.004 | 0.007 | CS | 0.00 | 0.00 | 0.00 | 0.00 | 100 | 0.00 | 0.00 |
| g_Phil1 | 0.009 | 0.014 | CSR | 0.00 | 0.00 | 0.00 | 0.00 | 0.00 | 0.40 | 99.60 |
| g_Aminivibrio | 0.050 | 0.067 | CSR | 0.00 | 0.00 | 0.00 | 0.00 | 0.00 | 0.00 | 100 |
| g_Micropruina | 0.065 | 0.084 | CSR | 0.00 | 0.00 | 0.00 | 0.00 | 0.00 | 0.00 | 100 |
| g_JAAZKI01 | 0.003 | 0.006 | S | 0.00 | 0.00 | 100 | 0.00 | 0.00 | 0.00 | 0.00 |
| g_SSC4 | 0.000* | 0.000* | C | 31.77 | 0.00 | 0.00 | 68.23 | 0.00 | 0.00 | 0.00 |
| g_DUNF01 | 0.100* | 0.125* | CSR | 0.00 | 0.00 | 0.00 | 0.00 | 0.00 | 0.00 | 100 |
| g_VBCG01 | 0.014 | 0.021 | S | 0.00 | 0.00 | 56.04 | 0.00 | 0.00 | 0.00 | 43.96 |
| g_RHKX01 | 0.000* | 0.000* | S | 0.00 | 0.00 | 98.90 | 0.00 | 0.00 | 0.00 | 1.10 |
| o__Thermomicrobiales | 0.343 | 0.370 | CSR | 0.00 | 0.00 | 0.00 | 0.00 | 0.00 | 0.00 | 100 |
| g_CAIXSE01 | 0.220 | 0.249 | CSR | 0.00 | 0.00 | 0.00 | 0.00 | 0.00 | 0.00 | 100 |
| f_CAG-74 | 0.038 | 0.052 | CSR | 0.00 | 0.00 | 0.00 | 0.00 | 0.00 | 0.00 | 100 |
| g_UBA4923 | 0.090 | 0.113 | CSR | 0.00 | 0.00 | 0.00 | 0.00 | 0.00 | 0.00 | 100 |
| g_JJ008 | 0.114 | 0.140 | CSR | 0.00 | 0.00 | 0.00 | 0.00 | 0.00 | 0.00 | 100 |
| f_UBA5550 | 0.000 | 0.001 | CSR | 0.00 | 0.00 | 0.00 | 0.00 | 0.00 | 0.00 | 100 |
| f_CFX8 | 0.004 | 0.007 | SR | 0.00 | 0.00 | 0.00 | 0.00 | 0.00 | 100 | 0.00 |
| f_Paludibacteraceae | 0.118 | 0.144 | CSR | 0.00 | 0.00 | 0.00 | 0.00 | 0.00 | 0.00 | 100 |
| g_OLB8 | 0.006 | 0.010 | CR | 0.00 | 0.00 | 0.00 | 100 | 0.00 | 0.00 | 0.00 |

| TAXA | Welch ANOVA p-value | BH[†] adjusted p-value | CSR‡ | Probability calculated by bootstrapping (%) | | | | | | |
|---|---|---|---|---|---|---|---|---|---|---|
| | | | | C[§] | R[§] | S[§] | CR[§] | CS[§] | SR[§] | CSR[§] |
| g_JAAYJV01 | 0.007 | 0.012 | CSR | 0.00 | 0.00 | 0.00 | 0.00 | 0.00 | 0.00 | 100 |
| g_Streptococcus | 0.002 | 0.005 | CS | 0.00 | 0.00 | 0.00 | 0.00 | 100 | 0.00 | 0.00 |
| g_HUN007 | 0.008 | 0.014 | CSR | 0.00 | 0.00 | 0.00 | 0.00 | 0.00 | 0.00 | 100 |
| f_SSC4 | 0.000* | 0.000* | SR | 0.00 | 0.00 | 1.00 | 0.00 | 0.00 | 99.00 | 0.00 |
| g_UBA1179 | 0.010* | 0.016* | CSR | 0.00 | 0.00 | 0.00 | 0.00 | 0.00 | 0.00 | 100 |
| g_JAAYFL01 | 0.054 | 0.071 | CSR | 0.00 | 0.00 | 0.00 | 0.00 | 0.00 | 0.00 | 100 |
| g_UBA1426 | 0.168 | 0.196 | CSR | 0.00 | 0.00 | 0.00 | 0.00 | 0.00 | 0.00 | 100 |
| g_IMCC26207 | 0.001 | 0.001 | S | 0.00 | 0.00 | 100 | 0.00 | 0.00 | 0.00 | 0.00 |
| g_UBA668 | 0.001 | 0.002 | C | 74.23 | 0.00 | 0.00 | 13.69 | 0.00 | 0.00 | 12.09 |
| g_DUPQ01 | 0.042 | 0.057 | S | 0.00 | 0.00 | 98.80 | 0.00 | 0.00 | 0.00 | 1.20 |
| f_CAG-826 | 0.042* | 0.057* | CSR | 0.00 | 0.00 | 0.00 | 0.00 | 0.00 | 0.00 | 100 |
| g_MWBF01 | 0.007 | 0.012 | CR | 0.00 | 0.00 | 0.00 | 100 | 0.00 | 0.00 | 0.00 |
| f_CAG-138 | 0.000 | 0.000 | R | 0.00 | 99.90 | 0.00 | 0.00 | 0.00 | 0.00 | 0.10 |
| f_Ilumatobacteraceae | 0.000* | 0.000* | CR | 0.00 | 0.00 | 0.00 | 100 | 0.00 | 0.00 | 0.00 |
| g_DUVQ01 | 0.000* | 0.000* | S | 0.00 | 0.00 | 100 | 0.00 | 0.00 | 0.00 | 0.00 |
| g_UBA5266 | 0.000* | 0.000* | CSR | 0.00 | 0.00 | 0.00 | 0.00 | 0.00 | 0.00 | 100 |
| f_Ruminococcaceae | 0.010 | 0.016 | S | 0.00 | 0.00 | 100 | 0.00 | 0.00 | 0.00 | 0.00 |
| g_CAIWTL01 | 0.000* | 0.000* | CR | 1.10 | 0.00 | 0.00 | 98.90 | 0.00 | 0.00 | 0.00 |
| g_Aestuariivirga | 0.000* | 0.000* | S | 0.00 | 0.00 | 53.55 | 0.00 | 46.45 | 0.00 | 0.00 |
| g_UBA8910 | 0.278 | 0.309 | CSR | 0.00 | 0.00 | 0.00 | 0.00 | 0.00 | 0.00 | 100 |
| g_CAG-170 | 0.000* | 0.000* | S | 0.00 | 0.00 | 100 | 0.00 | 0.00 | 0.00 | 0.00 |
| g_JAAEKA01 | 0.000* | 0.000* | CR | 0.00 | 0.00 | 0.00 | 100 | 0.00 | 0.00 | 0.00 |
| g_UBA10908 | 0.011 | 0.017 | CSR | 0.00 | 0.00 | 0.00 | 0.00 | 9.59 | 0.00 | 90.41 |
| g_MVYY01 | 0.000* | 0.000* | R | 0.00 | 100 | 0.00 | 0.00 | 0.00 | 0.00 | 0.00 |
| g_DTU049 | 0.000* | 0.000* | R | 0.00 | 100 | 0.00 | 0.00 | 0.00 | 0.00 | 0.00 |
| g_UBA1206 | 0.000* | 0.000* | CSR | 0.00 | 0.00 | 0.00 | 0.00 | 0.00 | 0.00 | 100 |
| g_RumEn-M2 | 0.000* | 0.000* | SR | 0.00 | 0.00 | 0.00 | 0.00 | 0.00 | 100 | 0.00 |
| g_JAAYTD01 | 0.000* | 0.000* | SR | 0.00 | 0.00 | 1.20 | 0.00 | 0.00 | 98.80 | 0.00 |
| g_Cloacibacterium | 0.000* | 0.000* | S | 0.00 | 0.00 | 99.60 | 0.00 | 0.00 | 0.40 | 0.00 |
| g_Promineofilum | 0.000* | 0.000* | CR | 0.00 | 0.50 | 0.00 | 99.50 | 0.00 | 0.00 | 0.00 |
| g_FeB-14 | 0.000* | 0.000* | CSR | 0.00 | 0.00 | 0.00 | 0.00 | 0.00 | 0.00 | 100 |
| g_Ancrocorticia | 0.000* | 0.000* | CSR | 0.00 | 0.00 | 0.00 | 0.00 | 0.00 | 0.00 | 100 |
| g_UBA1402 | 0.000* | 0.000* | S | 0.00 | 0.00 | 100 | 0.00 | 0.00 | 0.00 | 0.00 |
| g_JAAYNR01 | 0.000* | 0.000* | CSR | 0.00 | 0.00 | 0.00 | 0.00 | 0.00 | 0.00 | 100 |
| f_Pirellulaceae | 0.000* | 0.000* | CR | 0.20 | 0.00 | 0.00 | 99.80 | 0.00 | 0.00 | 0.00 |
| f_CAIOMD01 | 0.000* | 0.000* | S | 0.00 | 0.00 | 74.93 | 0.00 | 0.00 | 0.00 | 25.07 |
| g_SCN-69-89 | 0.000* | 0.000* | R | 0.00 | 100 | 0.00 | 0.00 | 0.00 | 0.00 | 0.00 |
| g_UBA412 | 0.000* | 0.000* | S | 0.00 | 0.00 | 100 | 0.00 | 0.00 | 0.00 | 0.00 |
| g_UBA4179 | 0.000* | 0.000* | SR | 0.00 | 0.10 | 1.30 | 0.00 | 0.00 | 97.50 | 1.10 |

| TAXA | Welch ANOVA p-value | BH† adjusted p-value | CSR‡ | Probability calculated by bootstrapping (%) | | | | | | |
|---|---|---|---|---|---|---|---|---|---|---|
| | | | | C§ | R§ | S§ | CR§ | CS§ | SR§ | CSR§ |
| f_Burkholderiaceae | 0.000* | 0.000* | CR | 1.70 | 0.00 | 0.00 | 98.30 | 0.00 | 0.00 | 0.00 |
| g_UBA2868 | 0.013* | 0.020* | CSR | 0.00 | 0.00 | 0.00 | 0.00 | 0.00 | 0.00 | 100 |
| g_GCA-2746885 | 0.000* | 0.000* | CS | 0.00 | 0.00 | 0.00 | 0.00 | 100 | 0.00 | 0.00 |
| g_UBA2224 | 0.000* | 0.000* | R | 0.00 | 100 | 0.00 | 0.00 | 0.00 | 0.00 | 0.00 |
| g_UBA4789 | 0.000* | 0.000* | SR | 0.00 | 0.00 | 1.30 | 0.00 | 0.00 | 82.72 | 15.98 |
| g_Accumulibacter | 0.000* | 0.000* | CR | 1.20 | 0.00 | 0.00 | 98.80 | 0.00 | 0.00 | 0.00 |
| g_JAABQG01 | 0.000* | 0.000* | R | 0.00 | 100 | 0.00 | 0.00 | 0.00 | 0.00 | 0.00 |
| g_Nocardioides | 0.000* | 0.000* | R | 0.00 | 100 | 0.00 | 0.00 | 0.00 | 0.00 | 0.00 |
| g_RXIX01 | 0.000* | 0.000* | R | 0.00 | 100 | 0.00 | 0.00 | 0.00 | 0.00 | 0.00 |
| g_RUG163 | 0.000* | 0.000* | S | 0.00 | 0.00 | 100 | 0.00 | 0.00 | 0.00 | 0.00 |
| g_GCA-013693735 | 0.000* | 0.000* | R | 0.00 | 100 | 0.00 | 0.00 | 0.00 | 0.00 | 0.00 |
| g_Novosphingobium | 0.000* | 0.000* | R | 0.00 | 100 | 0.00 | 0.00 | 0.00 | 0.00 | 0.00 |
| g_Petrimonas | 0.000* | 0.000* | S | 0.00 | 0.00 | 100 | 0.00 | 0.00 | 0.00 | 0.00 |
| f_DTU023 | 0.001* | 0.002* | C | 2.10 | 0.00 | 0.00 | 0.60 | 0.70 | 0.00 | 96.60 |
| o__Acetobacterales | 0.000* | 0.000* | S | 0.00 | 0.00 | 100 | 0.00 | 0.00 | 0.00 | 0.00 |
| f_Oscillospiraceae | 0.000* | 0.000* | R | 0.00 | 100 | 0.00 | 0.00 | 0.00 | 0.00 | 0.00 |
| g_UBA1384 | 0.013* | 0.020* | CSR | 0.00 | 0.40 | 0.00 | 0.40 | 0.00 | 0.10 | 99.10 |
| g_CAG-83 | 0.000* | 0.000* | S | 0.00 | 0.00 | 100 | 0.00 | 0.00 | 0.00 | 0.00 |
| f_Acetobacteraceae | 0.000* | 0.000* | R | 0.00 | 100 | 0.00 | 0.00 | 0.00 | 0.00 | 0.00 |
| f_Butyricicoccaceae | 0.000* | 0.000* | S | 0.00 | 0.00 | 100 | 0.00 | 0.00 | 0.00 | 0.00 |
| g_DTU032 | 0.000* | 0.000* | C | 100 | 0.00 | 0.00 | 0.00 | 0.00 | 0.00 | 0.00 |
| g_UBA1038 | 0.000* | 0.000* | R | 0.00 | 100 | 0.00 | 0.00 | 0.00 | 0.00 | 0.00 |
| g_DTU098 | 0.001* | 0.002* | S | 0.00 | 0.00 | 2.30 | 0.00 | 1.20 | 1.40 | 95.10 |
| g_UBA3907 | 0.000* | 0.000* | C | 100 | 0.00 | 0.00 | 0.00 | 0.00 | 0.00 | 0.00 |
| f_Syntrophomonadaceae | 0.000* | 0.000* | C | 100 | 0.00 | 0.00 | 0.00 | 0.00 | 0.00 | 0.00 |
| g_58-81 | 0.000* | 0.000* | R | 0.00 | 100 | 0.00 | 0.00 | 0.00 | 0.00 | 0.00 |
| f_JAAYNR01 | 0.000* | 0.000* | S | 0.00 | 0.00 | 100 | 0.00 | 0.00 | 0.00 | 0.00 |
| g_PNKZ01 | 0.000* | 0.000* | R | 0.00 | 100 | 0.00 | 0.00 | 0.00 | 0.00 | 0.00 |
| g_JACDBH01 | 0.000* | 0.000* | R | 0.00 | 100 | 0.00 | 0.00 | 0.00 | 0.00 | 0.00 |
| g_JAAYED01 | 0.000* | 0.000* | R | 0.00 | 100 | 0.00 | 0.00 | 0.00 | 0.00 | 0.00 |
| g_TTA-H9 | 0.000* | 0.000* | R | 0.00 | 100 | 0.00 | 0.00 | 0.00 | 0.00 | 0.00 |
| o__DTGP01 | 0.000* | 0.000* | R | 0.00 | 100 | 0.00 | 0.00 | 0.00 | 0.00 | 0.00 |

†: Benjamini-Hochberg

‡: CSR category assigned using the observed data based on Welch-ANOVA pairwise comparison between groups; C: competitor (no disturbance); r ruderal (low frequency of disturbance); R ruderal (intermediate frequency of disturbance); S: stress-tolerant (high frequency of disturbance)

§: Percentage of the genus gets assigned to each category calculated based on 1000 bootstrapping iteration

*: The variance was zero in at least one experimental group

**TABLE S3**. Summary statistics of multivariate analysis of variances on COG and CAZy annotation results using experimental group as explanatory variable

| Annotation | Term | Df* | Sums of squares | Means squares | F. Model | $R^2$ | P-value |
|---|---|---|---|---|---|---|---|
| COG | Experimental group | 3 | 0.0044 | 0.0015 | 52.048 | 0.9513 | 0.001 |
| | Residuals | 8 | 0.0002 | 0.0000 | | 0.0487 | |
| | Total | 11 | 0.0047 | | | 1.0000 | |
| CAZy | Experimental group | 3 | 0.0314 | 0.0105 | 35.453 | 0.0930 | 0.001 |
| | Residuals | 8 | 0.0024 | 0.0003 | | 0.0700 | |
| | Total | 11 | 0.0338 | | | 1.0000 | |

* Df: degrees of freedom

**TABLE S4**. Summary statistics of multivariate homogeneity of groups dispersions test on COG and CAZy annotations using experimental group as explanatory variable

| Annotation | Term | Df* | Sums of squares | Means squares | F. value | P-value |
|---|---|---|---|---|---|---|
| COG | Groups | 3 | $1.5316\times10^{-5}$ | $5.1055\times10^{-6}$ | 0.5376 | 0.6696 |
| | Residuals | 8 | $7.5972\times10^{-5}$ | $9.4965\times10^{-6}$ | | |
| CAZy | Groups | 3 | $1.5557\times10^{-5}$ | $5.1858\times10^{-6}$ | 0.2137 | 0.8842 |
| | Residuals | 8 | $1.9418\times10^{-4}$ | $2.4272\times10^{-5}$ | | |

* Df: degrees of freedom

**TABLE S5**. Summary of statistical analysis for classification of the COG category weighted counts under CSR framework. Table generated using MicroEcoTools R package (Neshat, Santillan and Wuertz, 2025).

| COG category | Description | Welch ANOVA p-value | CSR‡ | BH† adjusted p-value¶ | Probability calculated by bootstrapping (%) | | | | | | |
|---|---|---|---|---|---|---|---|---|---|---|---|
| | | | | | C§ | R§ | S§ | CR§ | CS§ | SR§ | CSR§ |
| A | RNA processing and modification | 0.006 | R | 0.015 | 1.10 | 17.28 | 0.00 | 29.07 | 0.00 | 0.00 | 52.55 |
| ADL | RNA processing and modification; Cell cycle control and mitosis; Replication and repair | 0.034 | CSR | 0.057 | 0.00 | 1.60 | 0.40 | 0.10 | 0.30 | 0.00 | 97.60 |
| AJ | RNA processing and modification; Translation | 0.302 | CSR | 0.341 | 0.40 | 0.00 | 0.00 | 0.00 | 0.20 | 0.00 | 99.40 |
| B | Chromatin Structure and dynamics | 0.000 | C | 0.003 | 87.11 | 0.00 | 0.00 | 0.50 | 0.10 | 0.00 | 12.29 |
| BDLTU | Chromatin Structure and dynamics; Cell cycle control and mitosis; Replication and repair; Signal Transduction; Intracellular trafficking and secretion; | 0.259 | CSR | 0.297 | 0.00 | 0.60 | 0.00 | 0.00 | 0.00 | 0.50 | 98.90 |
| BK | Chromatin Structure and dynamics; Transcription | 0.000 | R | 0.001 | 0.00 | 99.90 | 0.00 | 0.00 | 0.00 | 0.00 | 0.10 |
| BKQ | Chromatin Structure and dynamics; Transcription; Secondary Structure | 0.404 | CSR | 0.435 | 0.20 | 0.00 | 0.00 | 0.00 | 0.00 | 0.00 | 99.80 |
| BQ | Chromatin Structure and dynamics; Secondary Structure | 0.002 | R | 0.006 | 0.00 | 30.27 | 0.00 | 0.00 | 0.00 | 2.70 | 67.03 |
| C | Energy production and conversion | 0.005 | R | 0.013 | 0.00 | 94.41 | 0.00 | 0.00 | 0.00 | 0.00 | 5.59 |
| CE | Energy production and conversion; Amino Acid metabolism and transport | 0.014 | SR | 0.030 | 0.00 | 0.00 | 2.80 | 0.00 | 0.00 | 11.89 | 85.31 |
| CEH | Energy production and conversion; Amino Acid metabolism and transport; Coenzyme metabolism | 0.005 | C | 0.013 | 2.50 | 0.20 | 0.00 | 1.30 | 0.00 | 0.00 | 96.00 |
| CF | Energy production and conversion; Nucleotide metabolism and transport | 0.195 | CSR | 0.232 | 0.00 | 0.60 | 0.00 | 0.00 | 0.00 | 0.10 | 99.30 |
| CG | Energy production and conversion; Carbohydrate metabolism and transport | 0.022 | R | 0.041 | 0.00 | 21.58 | 0.00 | 0.20 | 0.00 | 0.00 | 78.22 |
| CGM | Energy production and conversion; Carbohydrate metabolism and transport; Cell wall/membrane/envelop biogenesis | 0.004 | CSR | 0.011 | 0.90 | 0.00 | 0.00 | 0.40 | 1.00 | 0.00 | 97.70 |
| CH | Energy production and conversion; Coenzyme metabolism | 0.001 | R | 0.003 | 0.10 | 90.31 | 0.00 | 0.30 | 0.00 | 0.00 | 9.29 |
| CHT | Energy production and conversion; Coenzyme metabolism; Signal Transduction | 0.002 | CR | 0.007 | 2.30 | 0.10 | 0.00 | 27.67 | 0.00 | 0.00 | 69.93 |
| CI | Energy production and conversion; Lipid metabolism | 0.068 | CSR | 0.099 | 0.20 | 0.50 | 0.00 | 2.30 | 0.00 | 0.00 | 97.00 |
| CJ | Energy production and conversion; Translation | 0.021 | CSR | 0.039 | 0.00 | 0.90 | 0.00 | 0.70 | 0.00 | 0.00 | 98.40 |
| CJQ | Energy production and conversion; Translation; Secondary Structure | 0.079 | CSR | 0.111 | 0.00 | 1.40 | 0.00 | 0.00 | 0.00 | 0.30 | 98.30 |
| CK | Energy production and conversion; Transcription | 0.434 | CSR | 0.464 | 0.00 | 0.60 | 0.00 | 0.00 | 0.20 | 0.00 | 99.20 |
| CM | Energy production and conversion; Cell wall/membrane/envelop biogenesis | 0.368 | CSR | 0.404 | 0.00 | 0.00 | 0.00 | 0.00 | 0.00 | 0.40 | 99.60 |
| CNT | Energy production and conversion; Cell motility; Signal Transduction | 0.002 | SR | 0.006 | 0.00 | 0.60 | 3.60 | 0.00 | 0.00 | 17.38 | 78.42 |

| COG category | Description | Welch ANOVA p-value | CSR[‡] | BH† adjusted p-value[¶] | Probability calculated by bootstrapping (%) | | | | | | |
|---|---|---|---|---|---|---|---|---|---|---|---|
| | | | | | C[§] | R[§] | S[§] | CR[§] | CS[§] | SR[§] | CSR[§] |
| CO | Energy production and conversion; Post-translational modification, protein turnover, chaperone functions | 0.000 | C | 0.002 | 95.80 | 0.00 | 0.00 | 4.20 | 0.00 | 0.00 | 0.00 |
| COP | Energy production and conversion; Post-translational modification, protein turnover, chaperone functions; Inorganic ion transport and metabolism | 0.001 | CR | 0.003 | 0.00 | 2.10 | 0.00 | 13.59 | 0.00 | 0.00 | 84.32 |
| COT | Energy production and conversion; Post-translational modification, protein turnover, chaperone functions; Signal Transduction | 0.688 | CSR | 0.692 | 0.10 | 0.10 | 0.10 | 0.00 | 0.00 | 0.40 | 99.30 |
| CP | Energy production and conversion; Inorganic ion transport and metabolism | 0.000 | R | 0.003 | 0.00 | 99.00 | 0.00 | 0.20 | 0.00 | 0.00 | 0.80 |
| CQ | Energy production and conversion; Secondary Structure | 0.150 | CSR | 0.192 | 0.00 | 0.00 | 0.00 | 0.00 | 0.40 | 0.00 | 99.60 |
| CT | Energy production and conversion; Signal Transduction | 0.042 | CSR | 0.066 | 0.30 | 0.30 | 0.00 | 4.50 | 0.00 | 0.00 | 94.91 |
| CU | Energy production and conversion; Intracellular trafficking and secretion | 0.007 | S | 0.017 | 0.00 | 0.00 | 26.27 | 0.00 | 0.10 | 2.10 | 71.53 |
| D | Cell cycle control and mitosis | 0.000 | R | 0.003 | 0.00 | 99.90 | 0.00 | 0.00 | 0.00 | 0.00 | 0.10 |
| DF | Cell cycle control and mitosis; Nucleotide metabolism and transport | 0.019 | CSR | 0.037 | 0.00 | 1.80 | 2.50 | 0.00 | 0.00 | 5.59 | 90.11 |
| DHM | Cell cycle control and mitosis; Coenzyme metabolism; Cell wall/membrane/envelop biogenesis | 0.027 | CSR | 0.046 | 1.60 | 0.00 | 0.00 | 0.00 | 1.70 | 0.00 | 96.70 |
| DJ | Cell cycle control and mitosis; Translation | 0.001 | R | 0.003 | 0.00 | 32.07 | 0.00 | 47.05 | 0.00 | 0.00 | 20.88 |
| DK | Cell cycle control and mitosis; Transcription | 0.054 | CSR | 0.082 | 0.90 | 3.10 | 0.00 | 2.90 | 0.00 | 0.00 | 93.11 |
| DL | Cell cycle control and mitosis; Replication and repair | 0.000 | R | 0.003 | 0.00 | 82.72 | 0.00 | 0.00 | 0.00 | 0.00 | 17.28 |
| DM | Cell cycle control and mitosis; Cell wall/membrane/envelop biogenesis | 0.000 | R | 0.003 | 0.00 | 54.85 | 0.00 | 35.36 | 0.00 | 0.00 | 9.79 |
| DMO | Cell cycle control and mitosis; Cell wall/membrane/envelop biogenesis; Post-translational modification, protein turnover, chaperone functions | 0.121 | CSR | 0.161 | 0.60 | 0.00 | 0.00 | 1.20 | 0.00 | 0.00 | 98.20 |
| DMUZ | Cell cycle control and mitosis; Cell wall/membrane/envelop biogenesis; Intracellular trafficking and secretion; Cytoskeleton | 0.002 | S | 0.005 | 0.00 | 0.00 | 19.68 | 0.00 | 1.70 | 0.00 | 78.62 |
| DMZ | Cell cycle control and mitosis; Cell wall/membrane/envelop biogenesis; Cytoskeleton | 0.096 | CSR | 0.131 | 0.00 | 0.00 | 0.00 | 0.00 | 0.50 | 0.00 | 99.50 |
| DN | Cell cycle control and mitosis; Cell motility | 0.022 | CSR | 0.041 | 0.00 | 0.60 | 0.90 | 0.00 | 0.00 | 5.39 | 93.11 |
| DO | Cell cycle control and mitosis; Post-translational modification, protein turnover, chaperone functions | 0.030 | CSR | 0.051 | 0.00 | 6.79 | 0.00 | 0.00 | 0.00 | 0.00 | 93.21 |
| DOZ | Cell cycle control and mitosis; Post-translational modification, protein turnover, chaperone functions; Cytoskeleton | 0.023 | R | 0.042 | 0.00 | 16.78 | 0.00 | 0.40 | 0.00 | 0.00 | 82.82 |
| DP | Cell cycle control and mitosis; Inorganic ion transport and metabolism | 0.004 | S | 0.012 | 0.00 | 0.00 | 15.58 | 0.00 | 0.90 | 0.30 | 83.22 |
| DT | Cell cycle control and mitosis; Signal Transduction | 0.004 | CR | 0.012 | 0.00 | 1.50 | 0.00 | 45.55 | 0.00 | 0.00 | 52.95 |
| DTZ | Cell cycle control and mitosis; Signal Transduction; Cytoskeleton | 0.139 | CSR | 0.182 | 0.00 | 0.10 | 0.00 | 0.00 | 0.00 | 0.00 | 99.90 |

| COG category | Description | Welch ANOVA p-value | CSR‡ | BH† adjusted p-value¶ | Probability calculated by bootstrapping (%) | | | | | | |
|---|---|---|---|---|---|---|---|---|---|---|---|
| | | | | | C§ | R§ | S§ | CR§ | CS§ | SR§ | CSR§ |
| DU | Cell cycle control and mitosis; Intracellular trafficking and secretion | 0.028 | CSR | 0.048 | 0.40 | 1.20 | 0.00 | 2.40 | 0.00 | 0.00 | 96.00 |
| DUZ | Cell cycle control and mitosis; Intracellular trafficking and secretion; Cytoskeleton | 0.005 | CR | 0.013 | 0.30 | 1.70 | 0.00 | 5.79 | 0.00 | 0.00 | 92.21 |
| DZ | Cell cycle control and mitosis; Cytoskeleton | 0.001 | C | 0.005 | 96.60 | 0.00 | 0.00 | 3.30 | 0.00 | 0.00 | 0.10 |
| E | Amino Acid metabolism and transport | 0.000 | CR | 0.003 | 0.00 | 0.00 | 0.00 | 88.71 | 0.00 | 0.00 | 11.29 |
| EF | Amino Acid metabolism and transport; Nucleotide metabolism and transport | 0.024 | CSR | 0.043 | 0.00 | 1.10 | 0.00 | 2.30 | 0.00 | 0.00 | 96.60 |
| EG | Amino Acid metabolism and transport; Carbohydrate metabolism and transport | 0.182 | CSR | 0.221 | 0.10 | 0.20 | 0.00 | 2.00 | 0.00 | 0.00 | 97.70 |
| EGH | Amino Acid metabolism and transport; Carbohydrate metabolism and transport; Coenzyme metabolism | 0.000 | CR | 0.003 | 8.39 | 0.20 | 0.00 | 57.34 | 0.00 | 0.00 | 34.07 |
| EGP | Amino Acid metabolism and transport; Carbohydrate metabolism and transport; Inorganic ion transport and metabolism | 0.117 | CSR | 0.156 | 0.00 | 0.80 | 0.00 | 0.00 | 0.00 | 0.00 | 99.20 |
| EH | Amino Acid metabolism and transport; Coenzyme metabolism | 0.026 | CR | 0.046 | 0.00 | 0.00 | 0.00 | 53.05 | 0.00 | 0.00 | 46.95 |
| EI | Amino Acid metabolism and transport; Lipid metabolism | 0.393 | CSR | 0.424 | 0.00 | 0.30 | 0.00 | 0.00 | 0.00 | 0.00 | 99.70 |
| EIP | Amino Acid metabolism and transport; Lipid metabolism; Inorganic ion transport and metabolism | 0.003 | S | 0.010 | 0.00 | 0.20 | 1.50 | 0.00 | 0.10 | 0.60 | 97.60 |
| EJ | Amino Acid metabolism and transport; Translation | 0.002 | R | 0.007 | 0.00 | 93.21 | 0.00 | 0.00 | 0.00 | 0.10 | 6.69 |
| EK | Amino Acid metabolism and transport; Transcription | 0.001 | R | 0.004 | 0.00 | 99.90 | 0.00 | 0.00 | 0.00 | 0.00 | 0.10 |
| ELM | Amino Acid metabolism and transport; Replication and repair; Cell wall/membrane/envelop biogenesis | 0.086 | CSR | 0.119 | 0.00 | 0.00 | 9.09 | 0.00 | 0.10 | 0.80 | 90.01 |
| EM | Amino Acid metabolism and transport; Cell wall/membrane/envelop biogenesis | 0.265 | CSR | 0.303 | 1.30 | 0.00 | 0.00 | 0.30 | 0.60 | 0.00 | 97.80 |
| EN | Amino Acid metabolism and transport; Cell motility | 0.323 | CSR | 0.359 | 0.00 | 0.00 | 0.00 | 0.00 | 0.00 | 0.00 | 100 |
| EP | Amino Acid metabolism and transport; Inorganic ion transport and metabolism | 0.129 | CSR | 0.171 | 0.00 | 0.80 | 0.00 | 0.00 | 0.00 | 0.00 | 99.20 |
| EQ | Amino Acid metabolism and transport; Secondary Structure | 0.005 | R | 0.013 | 0.00 | 20.98 | 0.00 | 0.40 | 0.00 | 0.00 | 78.62 |
| ET | Amino Acid metabolism and transport; Signal Transduction | 0.176 | CSR | 0.217 | 3.70 | 0.00 | 0.00 | 0.30 | 0.00 | 0.00 | 96.00 |
| EU | Amino Acid metabolism and transport; Intracellular trafficking and secretion | 0.010 | C | 0.022 | 33.47 | 0.00 | 0.00 | 0.00 | 0.30 | 0.00 | 66.23 |
| EUW | Amino Acid metabolism and transport; Intracellular trafficking and secretion | 0.667 | CSR | 0.677 | 0.00 | 0.10 | 0.30 | 0.20 | 0.30 | 0.10 | 99.00 |
| EV | Amino Acid metabolism and transport; Defence mechanisms | 0.027 | CSR | 0.046 | 0.00 | 5.99 | 0.00 | 0.00 | 0.00 | 0.10 | 93.91 |
| F | Nucleotide metabolism and transport | 0.000 | CR | 0.003 | 0.00 | 0.10 | 0.00 | 99.90 | 0.00 | 0.00 | 0.00 |
| FG | Nucleotide metabolism and transport; Carbohydrate metabolism and transport | 0.001 | R | 0.004 | 0.00 | 99.60 | 0.00 | 0.00 | 0.00 | 0.00 | 0.40 |

| COG category | Description | Welch ANOVA p-value | CSR[‡] | BH† adjusted p-value¶ | Probability calculated by bootstrapping (%) | | | | | | |
|---|---|---|---|---|---|---|---|---|---|---|---|
| | | | | | C[§] | R[§] | S[§] | CR[§] | CS[§] | SR[§] | CSR[§] |
| FGL | Nucleotide metabolism and transport; Carbohydrate metabolism and transport; Replication and repair | 0.520 | CSR | 0.536 | 0.00 | 0.10 | 0.00 | 0.00 | 0.00 | 0.00 | 99.90 |
| FGMQ | Nucleotide metabolism and transport; Carbohydrate metabolism and transport; Cell wall/membrane/envelop biogenesis; Secondary Structure | 0.019 | CSR | 0.037 | 4.40 | 0.00 | 0.00 | 0.00 | 1.00 | 0.00 | 94.61 |
| FH | Nucleotide metabolism and transport; Coenzyme metabolism | 0.000 | C | 0.002 | 83.72 | 0.00 | 0.00 | 12.99 | 0.00 | 0.00 | 3.30 |
| FIM | Nucleotide metabolism and transport; Lipid metabolism; Cell wall/membrane/envelop biogenesis | 0.577 | CSR | 0.591 | 0.00 | 0.00 | 0.40 | 0.00 | 0.10 | 0.10 | 99.40 |
| FJ | Nucleotide metabolism and transport; Translation | 0.000 | CR | 0.003 | 2.10 | 0.70 | 0.00 | 97.10 | 0.00 | 0.00 | 0.10 |
| FK | Nucleotide metabolism and transport; Transcription | 0.000 | CSR | 0.003 | 1.10 | 0.00 | 0.00 | 0.00 | 8.29 | 0.00 | 90.61 |
| FL | Nucleotide metabolism and transport; Replication and repair | 0.026 | CSR | 0.045 | 0.00 | 0.00 | 1.80 | 0.00 | 0.80 | 0.00 | 97.40 |
| FM | Nucleotide metabolism and transport; Cell wall/membrane/envelop biogenesis | 0.085 | CSR | 0.119 | 0.00 | 0.10 | 1.10 | 0.00 | 0.90 | 0.80 | 97.10 |
| FO | Nucleotide metabolism and transport; Post-translational modification, protein turnover, chaperone functions | 0.273 | CSR | 0.310 | 0.50 | 0.20 | 0.00 | 0.40 | 0.00 | 0.00 | 98.90 |
| FP | Nucleotide metabolism and transport; Inorganic ion transport and metabolism | 0.004 | R | 0.012 | 0.00 | 86.81 | 0.00 | 0.00 | 0.00 | 0.00 | 13.19 |
| FT | Nucleotide metabolism and transport; Signal Transduction | 0.166 | CSR | 0.208 | 0.00 | 1.20 | 0.10 | 0.00 | 0.00 | 0.00 | 98.70 |
| G | Carbohydrate metabolism and transport | 0.001 | CR | 0.003 | 0.00 | 27.07 | 0.00 | 25.87 | 0.00 | 0.00 | 47.05 |
| GH | Carbohydrate metabolism and transport; Coenzyme metabolism | 0.064 | CSR | 0.094 | 0.00 | 4.70 | 0.00 | 0.00 | 0.00 | 0.00 | 95.30 |
| GI | Carbohydrate metabolism and transport; Lipid metabolism | 0.064 | CSR | 0.094 | 0.10 | 0.20 | 0.00 | 0.00 | 0.40 | 0.00 | 99.30 |
| GIM | Carbohydrate metabolism and transport; Lipid metabolism; Cell wall/membrane/envelop biogenesis | 0.019 | CSR | 0.037 | 0.00 | 2.50 | 0.00 | 0.00 | 0.00 | 0.70 | 96.80 |
| GIQ | Carbohydrate metabolism and transport; Lipid metabolism; Secondary Structure | 0.071 | CSR | 0.101 | 0.00 | 0.00 | 1.50 | 0.00 | 0.00 | 0.40 | 98.10 |
| GJM | Carbohydrate metabolism and transport; Translation; Cell wall/membrane/envelop biogenesis | 0.140 | CSR | 0.182 | 0.00 | 1.40 | 0.00 | 0.30 | 0.00 | 0.00 | 98.30 |
| GK | Carbohydrate metabolism and transport; Transcription | 0.441 | CSR | 0.466 | 0.00 | 0.00 | 0.10 | 0.00 | 0.00 | 0.00 | 99.90 |
| GKM | Carbohydrate metabolism and transport; Transcription; Cell wall/membrane/envelop biogenesis | 0.257 | CSR | 0.297 | 0.00 | 0.00 | 0.10 | 0.00 | 0.00 | 0.00 | 99.90 |
| GKT | Carbohydrate metabolism and transport; Transcription; Signal Transduction | 0.019 | CSR | 0.037 | 0.00 | 2.80 | 0.20 | 0.00 | 0.00 | 5.69 | 91.31 |
| GL | Carbohydrate metabolism and transport; Replication and repair | 0.000 | R | 0.002 | 0.00 | 100 | 0.00 | 0.00 | 0.00 | 0.00 | 0.00 |
| GM | Carbohydrate metabolism and transport; Cell wall/membrane/envelop biogenesis | 0.001 | R | 0.005 | 0.00 | 57.94 | 0.00 | 22.58 | 0.00 | 0.00 | 19.48 |
| GMO | Carbohydrate metabolism and transport; Cell wall/membrane/envelop biogenesis; Post-translational modification, protein turnover, chaperone functions | 0.324 | CSR | 0.359 | 0.00 | 0.20 | 0.00 | 0.00 | 0.00 | 0.10 | 99.70 |

| COG category | Description | Welch ANOVA p-value | CSR‡ | BH† adjusted p-value¶ | Probability calculated by bootstrapping (%) | | | | | | |
|---|---|---|---|---|---|---|---|---|---|---|---|
| | | | | | C§ | R§ | S§ | CR§ | CS§ | SR§ | CSR§ |
| GN | Carbohydrate metabolism and transport; Cell motility | 0.092 | CSR | 0.126 | 4.30 | 0.00 | 0.00 | 0.00 | 0.20 | 0.00 | 95.50 |
| GNT | Carbohydrate metabolism and transport; Cell motility; Signal Transduction | 0.213 | CSR | 0.250 | 0.50 | 0.00 | 0.10 | 0.00 | 0.10 | 0.00 | 99.30 |
| GO | Carbohydrate metabolism and transport; Post-translational modification, protein turnover, chaperone functions | 0.003 | C | 0.009 | 86.91 | 0.00 | 0.00 | 0.00 | 0.00 | 0.00 | 13.09 |
| GP | Carbohydrate metabolism and transport; Inorganic ion transport and metabolism | 0.000 | CSR | 0.003 | 0.30 | 0.00 | 0.00 | 0.00 | 1.20 | 0.00 | 98.50 |
| GPT | Carbohydrate metabolism and transport; Inorganic ion transport and metabolism; Signal Transduction | 0.068 | CSR | 0.099 | 3.90 | 0.00 | 0.00 | 0.00 | 0.40 | 0.00 | 95.70 |
| GQ | Carbohydrate metabolism and transport; Secondary Structure | 0.001 | R | 0.003 | 0.00 | 89.21 | 0.00 | 0.00 | 0.00 | 0.00 | 10.79 |
| GT | Carbohydrate metabolism and transport; Signal Transduction | 0.000 | R | 0.002 | 0.30 | 86.61 | 0.00 | 6.29 | 0.00 | 0.00 | 6.79 |
| GV | Carbohydrate metabolism and transport; Defence mechanisms | 0.001 | S | 0.004 | 0.00 | 0.00 | 31.27 | 0.00 | 0.20 | 12.89 | 55.64 |
| H | Coenzyme metabolism | 0.007 | CR | 0.016 | 0.00 | 19.78 | 0.00 | 79.72 | 0.00 | 0.00 | 0.50 |
| HI | Coenzyme metabolism; Lipid metabolism | 0.382 | CSR | 0.415 | 0.00 | 0.00 | 0.00 | 0.00 | 0.10 | 0.00 | 99.90 |
| HJ | Coenzyme metabolism; Translation | 0.000 | R | 0.001 | 0.00 | 100 | 0.00 | 0.00 | 0.00 | 0.00 | 0.00 |
| HJK | Coenzyme metabolism; Translation; Transcription | 0.056 | CSR | 0.084 | 0.30 | 0.90 | 0.00 | 0.80 | 0.00 | 0.00 | 98.00 |
| HJM | Coenzyme metabolism; Translation; Cell wall/membrane/envelop biogenesis | 0.000 | CSR | 0.003 | 0.00 | 8.69 | 0.10 | 0.00 | 0.00 | 1.80 | 89.41 |
| HJO | Coenzyme metabolism; Translation; Post-translational modification, protein turnover, chaperone functions | 0.000 | CSR | 0.002 | 6.19 | 0.00 | 0.00 | 0.10 | 7.89 | 0.00 | 85.81 |
| HK | Coenzyme metabolism; Transcription | 0.002 | SR | 0.005 | 0.00 | 11.39 | 0.00 | 0.00 | 0.00 | 29.17 | 59.44 |
| HKT | Coenzyme metabolism; Transcription; Signal Transduction | 0.676 | CSR | 0.683 | 0.00 | 0.20 | 0.00 | 0.00 | 0.00 | 0.00 | 99.80 |
| HL | Coenzyme metabolism; Replication and repair | 0.505 | CSR | 0.526 | 0.00 | 0.00 | 0.20 | 0.00 | 0.10 | 0.00 | 99.70 |
| HM | Coenzyme metabolism; Cell wall/membrane/envelop biogenesis | 0.000 | C | 0.003 | 53.05 | 0.00 | 0.00 | 30.27 | 0.00 | 0.00 | 16.68 |
| HP | Coenzyme metabolism; Inorganic ion transport and metabolism | 0.001 | R | 0.003 | 0.00 | 92.41 | 0.00 | 0.00 | 0.00 | 0.70 | 6.89 |
| HQ | Coenzyme metabolism; Secondary Structure | 0.491 | CSR | 0.514 | 0.00 | 0.00 | 0.00 | 0.00 | 0.00 | 0.00 | 100 |
| I | Lipid metabolism | 0.005 | CSR | 0.013 | 0.00 | 0.80 | 0.00 | 0.40 | 0.00 | 0.00 | 98.80 |
| IJ | Lipid metabolism; Translation | 0.151 | CSR | 0.192 | 0.00 | 0.40 | 0.20 | 0.00 | 0.20 | 0.00 | 99.20 |
| IJM | Lipid metabolism; Translation; Cell wall/membrane/envelop biogenesis | 0.001 | CS | 0.004 | 0.10 | 0.00 | 0.30 | 0.00 | 3.70 | 0.10 | 95.80 |
| IK | Lipid metabolism; Transcription | 0.350 | CSR | 0.386 | 0.20 | 0.20 | 0.00 | 0.30 | 0.00 | 0.00 | 99.30 |
| IM | Lipid metabolism; Cell wall/membrane/envelop biogenesis | 0.001 | C | 0.003 | 81.22 | 0.00 | 0.00 | 0.00 | 7.29 | 0.00 | 11.49 |

| COG category | Description | Welch ANOVA p-value | CSR‡ | BH† adjusted p-value¶ | Probability calculated by bootstrapping (%) | | | | | | |
|---|---|---|---|---|---|---|---|---|---|---|---|
| | | | | | C§ | R§ | S§ | CR§ | CS§ | SR§ | CSR§ |
| IMQ | Lipid metabolism; Cell wall/membrane/envelop biogenesis; Secondary Structure | 0.015 | CSR | 0.031 | 1.10 | 1.10 | 0.00 | 3.70 | 0.00 | 0.00 | 94.11 |
| IMU | Lipid metabolism; Cell wall/membrane/envelop biogenesis; Intracellular trafficking and secretion | 0.046 | CSR | 0.072 | 0.00 | 0.00 | 0.60 | 0.00 | 0.30 | 0.10 | 99.00 |
| IN | Lipid metabolism; Cell motility | 0.178 | CSR | 0.218 | 0.90 | 0.10 | 0.10 | 0.10 | 0.20 | 0.00 | 98.60 |
| IO | Lipid metabolism; Post-translational modification, protein turnover, chaperone functions | 0.009 | CSR | 0.020 | 4.10 | 0.00 | 0.00 | 2.70 | 0.00 | 0.00 | 93.21 |
| IQ | Lipid metabolism; Secondary Structure | 0.208 | CSR | 0.246 | 0.00 | 0.00 | 0.00 | 0.00 | 0.00 | 0.00 | 100 |
| IT | Lipid metabolism; Signal Transduction | 0.001 | R | 0.004 | 6.49 | 24.98 | 0.00 | 22.18 | 0.00 | 0.00 | 46.35 |
| IU | Lipid metabolism; Intracellular trafficking and secretion | 0.001 | CR | 0.003 | 0.00 | 22.78 | 0.00 | 16.28 | 0.00 | 0.00 | 60.94 |
| J | Translation | 0.000 | R | 0.003 | 0.00 | 100 | 0.00 | 0.00 | 0.00 | 0.00 | 0.00 |
| JK | Translation; Transcription | 0.009 | CSR | 0.020 | 3.10 | 0.50 | 0.00 | 1.00 | 1.70 | 0.00 | 93.71 |
| JKL | Translation; Transcription; Replication and repair | 0.025 | CSR | 0.045 | 0.00 | 0.80 | 0.00 | 0.00 | 0.00 | 0.30 | 98.90 |
| JM | Translation; Cell wall/membrane/envelop biogenesis | 0.016 | CR | 0.032 | 8.39 | 0.10 | 0.00 | 4.80 | 0.20 | 0.00 | 86.51 |
| K | Transcription | 0.001 | CSR | 0.005 | 0.00 | 0.40 | 0.00 | 0.00 | 0.00 | 0.00 | 99.60 |
| KL | Transcription; Replication and repair | 0.005 | CR | 0.013 | 0.30 | 0.50 | 0.00 | 74.53 | 0.00 | 0.00 | 24.68 |
| KLT | Transcription; Replication and repair; Signal Transduction | 0.000 | R | 0.003 | 0.00 | 84.42 | 0.00 | 5.49 | 0.00 | 0.00 | 10.09 |
| KLTU | Transcription; Replication and repair; Signal Transduction; Intracellular trafficking and secretion | 0.001 | CR | 0.004 | 0.10 | 3.00 | 0.00 | 47.05 | 0.00 | 0.00 | 49.85 |
| KM | Transcription; Cell wall/membrane/envelop biogenesis | 0.236 | CSR | 0.276 | 1.10 | 0.00 | 0.00 | 0.10 | 0.10 | 0.00 | 98.70 |
| KMT | Transcription; Cell wall/membrane/envelop biogenesis; Signal Transduction | 0.150 | CSR | 0.192 | 1.90 | 0.00 | 0.00 | 0.40 | 0.10 | 0.00 | 97.60 |
| KNU | Transcription; Cell motility; Intracellular trafficking and secretion | 0.016 | CSR | 0.032 | 0.10 | 0.20 | 0.00 | 4.60 | 0.00 | 0.00 | 95.10 |
| KO | Transcription; Post-translational modification, protein turnover, chaperone functions | 0.737 | CSR | 0.737 | 0.00 | 0.00 | 0.00 | 0.00 | 0.00 | 0.00 | 100 |
| KOT | Transcription; Post-translational modification, protein turnover, chaperone functions; Signal Transduction | 0.037 | CSR | 0.061 | 0.00 | 1.20 | 0.00 | 0.00 | 0.00 | 0.10 | 98.70 |
| KP | Transcription; Inorganic ion transport and metabolism | 0.013 | CSR | 0.029 | 0.00 | 0.70 | 0.90 | 0.00 | 0.00 | 5.79 | 92.61 |
| KQ | Transcription; Secondary Structure | 0.001 | S | 0.004 | 0.00 | 0.00 | 76.62 | 0.00 | 0.60 | 1.80 | 20.98 |
| KT | Transcription; Signal Transduction | 0.000 | R | 0.002 | 0.00 | 95.20 | 0.00 | 0.30 | 0.00 | 0.00 | 4.50 |
| KTV | Transcription; Signal Transduction; Defence mechanisms | 0.594 | CSR | 0.606 | 0.00 | 0.00 | 0.00 | 0.00 | 0.00 | 0.00 | 100 |

| COG category | Description | Welch ANOVA p-value | CSR‡ | BH† adjusted p-value¶ | Probability calculated by bootstrapping (%) | | | | | | |
|---|---|---|---|---|---|---|---|---|---|---|---|
| | | | | | C§ | R§ | S§ | CR§ | CS§ | SR§ | CSR§ |
| KU | Transcription; Intracellular trafficking and secretion | 0.003 | SR | 0.009 | 0.00 | 6.49 | 0.00 | 0.00 | 0.00 | 15.48 | 78.02 |
| KV | Transcription; Defence mechanisms | 0.376 | CSR | 0.411 | 0.00 | 0.10 | 0.00 | 0.00 | 0.00 | 0.00 | 99.90 |
| L | Replication and repair | 0.001 | R | 0.004 | 0.00 | 97.90 | 0.00 | 2.10 | 0.00 | 0.00 | 0.00 |
| LM | Replication and repair; Cell wall/membrane/envelop biogenesis | 0.189 | CSR | 0.227 | 0.00 | 0.50 | 0.00 | 0.00 | 0.00 | 0.10 | 99.40 |
| LN | Replication and repair; Cell motility | 0.008 | CSR | 0.019 | 0.10 | 0.10 | 1.00 | 0.00 | 0.90 | 0.20 | 97.70 |
| LNU | Replication and repair; Cell motility; Intracellular trafficking and secretion | 0.307 | CSR | 0.345 | 0.00 | 0.00 | 0.00 | 0.00 | 0.00 | 0.10 | 99.90 |
| LO | Replication and repair; Post-translational modification, protein turnover, chaperone functions | 0.015 | CSR | 0.032 | 0.00 | 9.09 | 0.00 | 0.50 | 0.00 | 0.00 | 90.41 |
| LP | Replication and repair; Inorganic ion transport and metabolism | 0.001 | SR | 0.004 | 0.00 | 2.40 | 6.89 | 0.00 | 0.00 | 76.72 | 13.99 |
| LT | Replication and repair; Signal Transduction | 0.005 | R | 0.013 | 0.50 | 17.68 | 0.00 | 1.30 | 0.20 | 0.00 | 80.32 |
| LU | Replication and repair; Intracellular trafficking and secretion | 0.061 | CSR | 0.091 | 0.00 | 6.29 | 0.00 | 10.59 | 0.00 | 0.00 | 83.12 |
| LV | Replication and repair; Defence mechanisms | 0.000 | C | 0.003 | 100 | 0.00 | 0.00 | 0.00 | 0.00 | 0.00 | 0.00 |
| M | Cell wall/membrane/envelop biogenesis | 0.001 | CR | 0.003 | 0.00 | 0.00 | 0.00 | 100 | 0.00 | 0.00 | 0.00 |
| MN | Cell wall/membrane/envelop biogenesis; Cell motility | 0.040 | CSR | 0.064 | 0.90 | 0.00 | 0.00 | 0.00 | 2.90 | 0.00 | 96.20 |
| MNO | Cell wall/membrane/envelop biogenesis; Cell motility; Post-translational modification, protein turnover, chaperone functions | 0.048 | CSR | 0.075 | 0.00 | 0.10 | 0.00 | 0.50 | 0.00 | 0.00 | 99.40 |
| MNU | Cell wall/membrane/envelop biogenesis; Cell motility; Intracellular trafficking and secretion | 0.000 | C | 0.003 | 73.23 | 0.00 | 0.00 | 0.00 | 6.49 | 0.00 | 20.28 |
| MO | Cell wall/membrane/envelop biogenesis; post-translational modification, protein turnover, chaperone functions | 0.000 | C | 0.003 | 99.10 | 0.00 | 0.00 | 0.00 | 0.10 | 0.00 | 0.80 |
| MOQ | Cell wall/membrane/envelop biogenesis; Post-translational modification, protein turnover, chaperone functions; Secondary Structure | 0.078 | CSR | 0.111 | 2.60 | 0.10 | 0.00 | 1.50 | 0.00 | 0.00 | 95.80 |
| MP | Cell wall/membrane/envelop biogenesis; Inorganic ion transport and metabolism | 0.194 | CSR | 0.232 | 0.70 | 0.00 | 0.00 | 0.10 | 0.20 | 0.00 | 99.00 |
| MPU | Cell wall/membrane/envelop biogenesis; Inorganic ion transport and metabolism; Intracellular trafficking and secretion | 0.007 | CSR | 0.016 | 1.50 | 1.00 | 0.00 | 3.30 | 0.00 | 0.00 | 94.21 |
| MQ | Cell wall/membrane/envelop biogenesis; Secondary Structure | 0.018 | CSR | 0.036 | 6.79 | 0.00 | 0.00 | 0.00 | 6.59 | 0.00 | 86.61 |
| MQU | Cell wall/membrane/envelop biogenesis; Secondary Structure; Intracellular trafficking and secretion | 0.016 | SR | 0.032 | 0.00 | 1.80 | 1.00 | 0.00 | 0.00 | 12.89 | 84.32 |
| MT | Cell wall/membrane/envelop biogenesis; Signal Transduction | 0.006 | S | 0.016 | 0.00 | 0.00 | 88.41 | 0.00 | 0.00 | 0.00 | 11.59 |
| MU | Cell wall/membrane/envelop biogenesis; Intracellular trafficking and secretion | 0.001 | CR | 0.005 | 0.00 | 0.00 | 0.00 | 89.51 | 0.00 | 0.00 | 10.49 |
| MUW | Cell wall/membrane/envelop biogenesis; Intracellular trafficking and secretion | 0.105 | CSR | 0.142 | 2.70 | 0.00 | 0.00 | 0.40 | 0.20 | 0.00 | 96.70 |

| COG category | Description | Welch ANOVA p-value | CSR‡ | BH† adjusted p-value¶ | Probability calculated by bootstrapping (%) | | | | | | |
|---|---|---|---|---|---|---|---|---|---|---|---|
| | | | | | C§ | R§ | S§ | CR§ | CS§ | SR§ | CSR§ |
| MV | Cell wall/membrane/envelop biogenesis; Defence mechanisms | 0.010 | CSR | 0.022 | 0.00 | 1.60 | 0.00 | 2.90 | 0.00 | 0.00 | 95.50 |
| N | Cell motility | 0.003 | R | 0.008 | 0.00 | 100 | 0.00 | 0.00 | 0.00 | 0.00 | 0.00 |
| NO | Cell motility; Post-translational modification, protein turnover, chaperone functions | 0.039 | CSR | 0.063 | 0.10 | 0.30 | 0.00 | 2.30 | 0.00 | 0.00 | 97.30 |
| NOT | Cell motility; Post-translational modification, protein turnover, chaperone functions; Signal Transduction | 0.439 | CSR | 0.466 | 0.00 | 0.00 | 0.00 | 0.00 | 0.00 | 0.00 | 100 |
| NOU | Cell motility; Post-translational modification, protein turnover, chaperone functions; Intracellular trafficking and secretion | 0.003 | R | 0.008 | 0.00 | 95.50 | 0.00 | 0.90 | 0.00 | 0.00 | 3.60 |
| NPTU | Cell motility; Inorganic ion transport and metabolism; Signal Transduction; Intracellular trafficking and secretion | 0.000 | C | 0.002 | 99.90 | 0.00 | 0.00 | 0.00 | 0.00 | 0.00 | 0.10 |
| NPU | Cell motility; Inorganic ion transport and metabolism; Intracellular trafficking and secretion | 0.105 | CSR | 0.142 | 0.00 | 0.50 | 0.50 | 0.10 | 0.00 | 0.30 | 98.60 |
| NQ | Cell motility; Secondary Structure | 0.169 | CSR | 0.210 | 0.00 | 0.00 | 0.20 | 0.00 | 0.00 | 0.00 | 99.80 |
| NT | Cell motility; Signal Transduction | 0.001 | R | 0.003 | 0.00 | 96.90 | 0.00 | 0.00 | 0.00 | 0.00 | 3.10 |
| NTU | Cell motility; Signal Transduction; Intracellular trafficking and secretion | 0.040 | CSR | 0.064 | 0.00 | 0.30 | 0.00 | 0.00 | 0.00 | 0.60 | 99.10 |
| NU | Cell motility; Intracellular trafficking and secretion | 0.001 | R | 0.003 | 0.00 | 100 | 0.00 | 0.00 | 0.00 | 0.00 | 0.00 |
| NUV | Cell motility; Intracellular trafficking and secretion; Defence mechanisms | 0.148 | CSR | 0.192 | 0.00 | 1.50 | 0.00 | 0.00 | 0.00 | 0.00 | 98.50 |
| O | Post-translational modification, protein turnover, chaperone functions | 0.000 | CR | 0.003 | 0.00 | 0.20 | 0.00 | 99.80 | 0.00 | 0.00 | 0.00 |
| OP | Post-translational modification, protein turnover, chaperone functions; Inorganic ion transport and metabolism | 0.041 | CSR | 0.065 | 0.00 | 0.30 | 1.50 | 0.00 | 0.00 | 1.20 | 97.00 |
| OQ | Post-translational modification, protein turnover, chaperone functions; Secondary Structure | 0.164 | CSR | 0.207 | 0.80 | 0.00 | 0.00 | 0.70 | 0.00 | 0.00 | 98.50 |
| OT | Post-translational modification, protein turnover, chaperone functions; Signal Transduction | 0.051 | CSR | 0.078 | 0.00 | 0.00 | 0.00 | 0.00 | 0.00 | 1.30 | 98.70 |
| OU | Post-translational modification, protein turnover, chaperone functions; Intracellular trafficking and secretion | 0.000 | R | 0.002 | 0.00 | 100 | 0.00 | 0.00 | 0.00 | 0.00 | 0.00 |
| OW | Post-translational modification, protein turnover, chaperone functions | 0.037 | CSR | 0.061 | 0.00 | 23.98 | 0.00 | 0.10 | 0.00 | 0.00 | 75.92 |
| P | Inorganic ion transport and metabolism | 0.008 | CSR | 0.019 | 0.00 | 14.69 | 0.00 | 13.39 | 0.00 | 0.00 | 71.93 |
| PQ | Inorganic ion transport and metabolism; Secondary Structure | 0.027 | R | 0.046 | 0.00 | 12.59 | 0.00 | 0.00 | 0.00 | 0.10 | 87.31 |
| PT | Inorganic ion transport and metabolism; Signal Transduction | 0.183 | CSR | 0.222 | 0.00 | 0.00 | 0.00 | 0.00 | 0.00 | 0.00 | 100 |
| PU | Inorganic ion transport and metabolism; Intracellular trafficking and secretion | 0.034 | CR | 0.057 | 0.80 | 0.50 | 0.00 | 5.69 | 0.00 | 0.00 | 93.01 |
| PV | Inorganic ion transport and metabolism; Defence mechanisms | 0.070 | CSR | 0.100 | 0.70 | 0.10 | 0.00 | 0.20 | 0.00 | 0.00 | 99.00 |

| COG category | Description | Welch ANOVA p-value | CSR[‡] | BH† adjusted p-value[¶] | Probability calculated by bootstrapping (%) | | | | | | |
|---|---|---|---|---|---|---|---|---|---|---|---|
| | | | | | C[§] | R[§] | S[§] | CR[§] | CS[§] | SR[§] | CSR[§] |
| Q | Secondary Structure | 0.001 | CR | 0.004 | 0.00 | 1.00 | 0.00 | 81.22 | 0.00 | 0.00 | 17.78 |
| QT | Secondary Structure; Signal Transduction | 0.512 | CSR | 0.531 | 0.00 | 0.00 | 0.00 | 0.00 | 0.00 | 0.00 | 100 |
| QU | Secondary Structure; Intracellular trafficking and secretion | 0.002 | CSR | 0.005 | 0.90 | 3.70 | 0.00 | 26.27 | 0.00 | 0.00 | 69.13 |
| QUW | Secondary Structure; Intracellular trafficking and secretion | 0.168 | CSR | 0.209 | 0.80 | 0.60 | 0.00 | 0.20 | 0.00 | 0.10 | 98.30 |
| S | Function Unknown | 0.001 | CR | 0.003 | 0.00 | 0.00 | 0.00 | 100 | 0.00 | 0.00 | 0.00 |
| T | Signal Transduction | 0.001 | R | 0.005 | 0.00 | 99.90 | 0.00 | 0.00 | 0.00 | 0.00 | 0.10 |
| TU | Signal Transduction; Intracellular trafficking and secretion | 0.052 | CSR | 0.080 | 0.40 | 13.99 | 0.00 | 1.20 | 0.00 | 0.00 | 84.42 |
| TV | Signal Transduction; Defence mechanisms | 0.246 | CSR | 0.286 | 0.40 | 0.10 | 0.00 | 0.20 | 0.00 | 0.00 | 99.30 |
| TZ | Signal Transduction; Cytoskeleton | 0.476 | CSR | 0.501 | 0.00 | 0.00 | 0.00 | 0.00 | 0.00 | 0.00 | 100 |
| U | Intracellular trafficking and secretion | 0.001 | R | 0.003 | 0.00 | 91.11 | 0.00 | 6.19 | 0.00 | 0.00 | 2.70 |
| UW | Intracellular trafficking and secretion | 0.016 | CR | 0.032 | 3.90 | 0.00 | 0.00 | 23.48 | 0.00 | 0.00 | 72.63 |
| V | Defence mechanisms | 0.000 | CR | 0.002 | 0.00 | 0.10 | 0.00 | 99.90 | 0.00 | 0.00 | 0.00 |
| Z | Cytoskeleton | 0.015 | CSR | 0.032 | 2.50 | 0.00 | 0.00 | 3.00 | 0.00 | 0.00 | 94.51 |

§: Percentage of the genus gets assigned to each category calculated based on 1000 bootstrapping iteration

‡: CSR category assigned using the observed data based on Welch-ANOVA pairwise comparison between groups; C: competitor (no disturbance); r ruderal (low frequency of disturbance); R ruderal (intermediate frequency of disturbance); S: stress-tolerant (high frequency of disturbance)

†: Benjamini-Hochberg

**TABLE S6**. Summary of statistical analysis for classification of the CAZy category weighted counts under CSR framework. Table generated using MicroEcoTools R package (Neshat, Santillan and Wuertz, 2025).

| CAZy category‡ | Welch ANOVA p-value | BH† adjusted p-value | Cr§ | CR§ | CS§ | rR§ | rS§ | RS§ | CSR |
|---|---|---|---|---|---|---|---|---|---|
| GH15 | 0.000 | 0.000 | C | C | C | r | S | S | C |
| GH29 | 0.010 | 0.022 | C | C | C | N | N | N | C |
| GH31 | 0.029 | 0.052 | C | C | C | N | N | N | C |
| GH37,GH65 | 0.001 | 0.006 | C | C | C | N | S | N | C |
| GH65,GT3,GT35 | 0.001 | 0.004 | C | C | C | N | N | N | C |
| GH66 | 0.002 | 0.006 | C | C | C | N | N | R | C |
| GT5 | 0.003 | 0.011 | C | C | C | N | N | N | C |
| GT9 | 0.002 | 0.006 | C | C | C | R | N | N | C |
| CBM50 | 0.001 | 0.006 | N | C | C | r | R | R | CR |
| GH105,GH28 | 0.002 | 0.008 | C | N | C | R | N | R | CR |
| GH31,GH77 | 0.001 | 0.006 | C | N | C | R | R | R | CR |
| GH39 | 0.013 | 0.026 | N | N | C | N | R | R | CR |
| GH5,GH9 | 0.005 | 0.014 | N | C | C | N | R | R | CR |
| GT1 | 0.001 | 0.006 | N | N | C | R | R | R | CR |
| GT30 | 0.000 | 0.002 | N | C | C | r | R | R | CR |
| GT51 | 0.005 | 0.014 | N | N | C | N | R | R | CR |
| AA10,CBM73 | 0.003 | 0.010 | N | N | N | N | S | S | CSR |
| CBM18,GH19 | 0.049 | 0.080 | N | N | N | N | N | N | CSR |
| CBM20,GH13 | 0.459 | 0.518 | N | N | N | N | N | N | CSR |
| CBM48,GH13 | 0.007 | 0.018 | N | N | N | N | N | R | CSR |
| CBM5,GH18 | 0.132 | 0.180 | N | N | N | N | N | N | CSR |
| CBM6,GH43,GH5 | 0.569 | 0.595 | N | N | N | N | N | N | CSR |
| CE1 | 0.273 | 0.337 | N | N | N | N | N | N | CSR |
| GH1,GH3 | 0.308 | 0.361 | N | N | N | N | N | N | CSR |
| GH101 | 0.009 | 0.020 | N | N | N | N | N | N | CSR |
| GH101,GH13,GH29,GH31 | 0.009 | 0.020 | R | N | N | N | N | N | CSR |
| GH101,GH29 | 0.008 | 0.020 | N | N | N | R | N | R | CSR |
| GH102 | 0.053 | 0.083 | N | N | N | N | N | N | CSR |
| GH105 | 0.007 | 0.018 | N | N | N | r | N | N | CSR |
| GH116 | 0.632 | 0.649 | N | N | N | N | N | N | CSR |
| GH13,GH26,GH29 | 0.063 | 0.094 | N | N | N | N | N | N | CSR |
| GH13,GH31 | 0.021 | 0.039 | N | N | N | N | N | N | CSR |
| GH13,GH31,GH77 | 0.159 | 0.204 | N | N | N | N | N | N | CSR |
| GH13,GH5,GH9 | 0.431 | 0.493 | N | N | N | N | N | N | CSR |
| GH13,GT5 | 0.043 | 0.071 | N | N | N | N | N | N | CSR |
| GH16 | 0.275 | 0.337 | N | N | N | N | N | N | CSR |
| GH19 | 0.081 | 0.117 | N | N | N | N | N | N | CSR |
| GH20,GH33 | 0.681 | 0.693 | N | N | N | N | N | N | CSR |

| CAZy category[‡] | Welch ANOVA p-value | BH[†] adjusted p-value | Cr[§] | CR[§] | CS[§] | rR[§] | rS[§] | RS[§] | CSR |
|---|---|---|---|---|---|---|---|---|---|
| GH26 | 0.013 | 0.026 | N | N | N | N | N | N | CSR |
| GH26,GH29 | 0.053 | 0.083 | N | N | N | N | N | N | CSR |
| GH26,GH5,GH9 | 0.165 | 0.208 | N | N | N | N | N | N | CSR |
| GH3 | 0.001 | 0.004 | R | C | N | r | N | S | CSR |
| GH32 | 0.064 | 0.094 | N | N | N | N | N | N | CSR |
| GH32,GH8 | 0.974 | 0.974 | N | N | N | N | N | N | CSR |
| GH33,GH5,GH9 | 0.074 | 0.108 | N | N | N | N | N | N | CSR |
| GH37 | 0.113 | 0.159 | N | N | N | N | N | N | CSR |
| GH4,GT4 | 0.056 | 0.086 | N | N | N | N | N | N | CSR |
| GH43,GH51 | 0.052 | 0.083 | N | N | N | N | N | N | CSR |
| GH48,GH5,GH9 | 0.062 | 0.094 | N | N | N | N | N | N | CSR |
| GH5 | 0.295 | 0.350 | N | N | N | N | N | N | CSR |
| GH5,GH9,GT2 | 0.238 | 0.297 | N | N | N | N | N | N | CSR |
| GH5,GH9,GT39,GT87 | 0.133 | 0.180 | N | N | N | N | N | N | CSR |
| GH51 | 0.530 | 0.559 | N | N | N | N | N | N | CSR |
| GH6,GH82 | 0.017 | 0.033 | N | N | N | N | N | N | CSR |
| GH65 | 0.014 | 0.028 | N | N | N | r | N | N | CSR |
| GH65,GT35 | 0.286 | 0.343 | N | N | N | N | N | N | CSR |
| GH77 | 0.050 | 0.081 | N | N | N | r | N | N | CSR |
| GH8 | 0.279 | 0.338 | N | N | N | N | N | N | CSR |
| GH84 | 0.423 | 0.492 | N | N | N | N | N | N | CSR |
| GH88 | 0.623 | 0.646 | N | N | N | N | N | N | CSR |
| GH94 | 0.138 | 0.182 | N | N | N | N | N | N | CSR |
| GH94,GT36 | 0.470 | 0.524 | N | N | N | N | N | N | CSR |
| GH94,GT36,GT84 | 0.146 | 0.189 | N | N | N | N | N | N | CSR |
| GH94,GT84 | 0.040 | 0.069 | N | N | N | N | N | N | CSR |
| GH95 | 0.004 | 0.014 | N | C | N | r | N | N | CSR |
| GT10 | 0.713 | 0.720 | N | N | N | N | N | N | CSR |
| GT11 | 0.495 | 0.537 | N | N | N | N | N | N | CSR |
| GT19 | 0.041 | 0.069 | N | N | N | N | N | N | CSR |
| GT2,GT4 | 0.028 | 0.051 | N | R | N | R | N | N | CSR |
| GT20 | 0.485 | 0.531 | N | N | N | N | N | N | CSR |
| GT21 | 0.484 | 0.531 | N | N | N | N | N | N | CSR |
| GT25 | 0.090 | 0.128 | N | N | N | N | N | N | CSR |
| GT3 | 0.433 | 0.493 | N | N | N | N | N | N | CSR |
| GT3,GT35 | 0.136 | 0.182 | N | N | N | N | N | N | CSR |
| GT36 | 0.005 | 0.014 | R | N | N | N | N | N | CSR |
| GT4 | 0.009 | 0.020 | N | N | N | N | N | N | CSR |
| GT56 | 0.012 | 0.025 | N | N | N | N | N | N | CSR |
| GT6 | 0.502 | 0.540 | N | N | N | N | N | N | CSR |
| GT66 | 0.129 | 0.179 | N | N | N | N | N | N | CSR |

| CAZy category[‡] | Welch ANOVA p-value | BH[†] adjusted p-value | Cr[§] | CR[§] | CS[§] | rR[§] | rS[§] | RS[§] | CSR |
|---|---|---|---|---|---|---|---|---|---|
| GT81 | 0.146 | 0.189 | N | N | N | N | N | N | CSR |
| GT87 | 0.030 | 0.053 | N | N | N | N | N | N | CSR |
| PL11 | 0.527 | 0.559 | N | N | N | N | N | N | CSR |
| PL12 | 0.006 | 0.017 | C | N | N | R | S | N | CSR |
| PL8 | 0.009 | 0.020 | N | R | N | R | N | N | CSR |
| CBM48,GH13,GH77 | 0.000 | 0.002 | R | R | C | N | R | R | R |
| CE10 | 0.000 | 0.002 | C | R | C | R | N | R | R |
| GH103 | 0.006 | 0.016 | N | R | N | R | N | R | R |
| GH121 | 0.011 | 0.024 | N | R | N | R | N | R | R |
| GH13 | 0.001 | 0.005 | R | R | N | N | R | R | R |
| GH18 | 0.000 | 0.002 | C | R | C | R | N | R | R |
| GH18,GH5,GH9 | 0.000 | 0.002 | N | R | C | R | N | R | R |
| GH20 | 0.001 | 0.006 | C | R | C | R | N | R | R |
| GH23 | 0.000 | 0.002 | R | R | N | N | R | R | R |
| GH30 | 0.008 | 0.020 | R | N | N | N | R | N | R |
| GH33 | 0.003 | 0.010 | R | R | C | r | R | R | R |
| GH38 | 0.001 | 0.006 | R | R | C | N | R | R | R |
| GH43 | 0.000 | 0.002 | R | C | C | r | R | N | R |
| GH57 | 0.002 | 0.006 | R | N | C | N | R | R | R |
| GH6 | 0.000 | 0.002 | R | R | N | R | N | R | R |
| GH68 | 0.003 | 0.010 | R | C | N | r | R | N | R |
| GT2 | 0.000 | 0.002 | R | R | C | R | R | R | R |
| GT26 | 0.000 | 0.002 | N | R | S | N | N | R | R |
| GT28 | 0.001 | 0.006 | R | N | C | r | R | R | R |
| GT35 | 0.000 | 0.003 | N | R | C | R | R | R | R |
| GT39 | 0.005 | 0.014 | R | R | N | N | R | R | R |
| GT89 | 0.000 | 0.003 | C | R | C | R | N | R | R |
| CBM6,GH43 | 0.004 | 0.012 | N | C | S | r | S | S | S |
| PL21 | 0.019 | 0.036 | N | N | S | N | S | S | S |
| CBM48,GH13,GH31 | 0.001 | 0.006 | R | R | S | N | N | N | SR |
| GH13,GT4,GT5 | 0.002 | 0.006 | R | N | S | r | N | S | SR |
| GH130 | 0.009 | 0.020 | R | R | S | N | N | N | SR |
| GH29,GH43 | 0.000 | 0.002 | N | R | S | R | S | N | SR |
| GH31,GH97 | 0.022 | 0.039 | N | R | S | R | S | N | SR |
| GH63 | 0.000 | 0.004 | R | N | S | r | N | S | SR |
| GT2,GT81 | 0.006 | 0.016 | R | N | S | r | N | S | SR |

§: Pairwise comparison between groups C: competitor (no disturbance); r ruderal (low frequency of disturbance); R ruderal (intermediate frequency of disturbance); S: stress-tolerant (high frequency of disturbance)

†: Benjamini-Hochberg

‡: GH: Glycoside Hydrolases (hydrolysis and/or rearrangement of glycosidic bonds); GT: Glycosyl Transferases (formation of glycosidic bonds); PL: Polysaccharide Lyases (non-

hydrolytic cleavage of glycosidic bonds); CE: Carbohydrate Esterases (hydrolysis of carbohydrate esters); AA: Auxiliary Activities (redox enzymes that act in conjunction with CAZymes); CBM: Carbohydrate-Binding Modules (adhesion to carbohydrates)
